# Physiological diversity and adaptation of Rhizaria revealed by phylogenomics and comparative transcriptomics

**DOI:** 10.1101/2025.10.20.683504

**Authors:** Jule Freudenthal, Michael Bonkowski, Nele-Joelle Maria Engelmann, Martin Schlegel, Kenneth Dumack

## Abstract

Protists are vastly diverse, forming over 20 supergroups of the eukaryotic diversity and fulfilling plentiful functions. Rhizaria is a widespread and highly abundant supergroup comprising important parasites and a huge diversity of free-living heterotrophic predators. Despite the diversity and biogeochemical importance of Rhizaria, our understanding of their physiology and metabolic capabilities remains limited, mainly due to a general lack of data and bioinformatic tools for cross-species comparisons of physiological traits. In this study, we assembled a total of 15 transcriptomes of the parasite-related bacterivorous *Rhogostoma* and their eukaryvorous relatives. By phylogenomic analyses and whole transcriptome comparison, we established an evolutionary framework to which we relate physiological traits. The morphologically highly similar *Rhogostoma* strains branch in two distinct clusters differing in orthogroups and gene expression patterns related to cell adhesion and biofilm formation. Furthermore, we reveal considerable intra-genus variation in amino acid and lipid metabolism, which might be explained by an ancient streamlining through gradual specialization to parasitism, bearing the potential for subsequent metabolic radiation. We conclude that even closely related and morphologically similar species in Rhizaria may differ distinctly in their functional repertoire. With the here established and showcased analyses, we create a basis for future characterization of the physiological traits of microeukaryotes.

## 1. Introduction

The Rhizaria are among the least understood but most diverse microbial eukaryotes. Rhizaria exhibits a high morphological and ecological diversity, including (1) Foraminifera, Radiolaria, and Phaeodaria, which are important oceanic predators; (2) phototrophic taxa (algae), such as *Paulinella chromatophora* and Chlorarachniophytes, which are crucial for our understanding of the evolution of photosynthesis; and (3) numerous parasitic species, such as *Plasmodiophora brassicae*, which causes about 15% of cabbage yields losses worldwide (Cavalier-Smith et al., 2018; Dumack et al., 2020; Nakamura and Suzuki, 2015; Neuhauser et al., 2010; Nowack, 2014). In addition, Rhizaria includes a variety of free-living and heterotrophic species, many of which exhibit an enormous species richness and genetic diversity (Bass et al., 2009; Flues et al., 2018; Howe et al., 2009).

Thecofilosea is one of the most abundant Rhizaria phyla across ecosystems, for example, in soils and the polar oceans, as shown by global surveys (Oliverio et al., 2020; Sommeria-Klein et al., 2021). Thecofilosea comprise a remarkable ecological and morphological diversity, including bacterivores, eukaryvores, and specialized, parasite-like predators of algae (Dumack et al., 2020). Among the Thecofilosea, free-living and heterotrophic taxa of the genus *Rhogostoma* (Rhogostomidae) are most abundant in terrestrial ecosystems (Öztoprak et al., 2020; Walden et al., 2021). *Rhogostoma* viciously preys on bacteria, whereas most of its relatives feed on eukaryotes (Seppey et al., 2017). Furthermore, *Rhogostoma* exhibits remarkable 18S marker gene variation but a highly conserved morphology, indicating ongoing cryptic speciation with a hidden diversity that remains to be fully uncovered (Öztoprak et al., 2020).

Despite the widespread occurrence and high abundance of Rhizaria, and *Rhogostoma* in particular, we still lack a well-supported phylogeny of the Rhogostomidae as single-or few-gene phylogenies, in contrast to many other microbial taxa, do not resolve most interspecific phylogenetic relationships (Cavalier-Smith et al., 2018). Furthermore, the transcriptome representation of Rhizaria is scarce (Sibbald and Archibald, 2017), and their physiology and functional diversity remain to be explored. With modern high-throughput sequencing techniques and bioinformatics tools, it is now feasible to explore the physiological traits of individual species in an evolutionary context, which in turn will shed light on their potential ecological impact (Gerbracht et al., 2022; Ribeiro et al., 2020).

In this study, we explore the phylogenetic relationships and physiological traits of Thecofilosea. We present a compiled data set of 12 novel *Rhogostoma* transcriptomes, two Tectofilosida transcriptomes, and one Ebriida sp. transcriptome. Using a multi-gene phylogenomic approach, we resolve the phylogenetic backbone of Thecofilosea, in particular of *Rhogostoma*. Additionally, we perform whole transcriptome comparisons, along with functional annotation and enrichment analyses, to explore the inter- and intraspecific diversity in the physiological traits of the Thecofilosea in an evolutionary context.

## 2. Methods

### 2.1. RNA Extraction

We extracted total RNA of monoclonal cultures of 11 *Rhogostoma* strains previously described and cultured by Öztoprak et al. (2020), Pohl et al. (2021), and Martínez Rendón et al. (2025), along with *Katarium polorum* (Solbach et al., 2025). Additionally, one *Rhogostoma* strain was isolated and cultured from a sample originating from the Leipzig floodplain forest (51.3657 N, 12.3094 E) in Germany by isolating single cells using sterile glass micropipettes and culturing them in cell culture flasks (SARSTEDT AG & Co. KG T25; Nümbrecht, Germany) with wheat grass (WG)-medium, at temperatures ranging from 4 to 21°C.

For each RNA extraction from the cultures, cells were detached from the bottles by vigorous shaking and/or thorough scraping with a sterile cell scraper. Subsequently, the medium was filtered with a filter pore size of 3 μm (cellulose nitrate membranes, Whatman™, Buckinghamshire, United Kingdom) until the filter was completely covered in cells. The filter was transferred to a 1.5 ml tube (SARSTEDT AG & Co. KG, Nümbrecht, Germany) containing 1 ml ice-cold Sørensen buffer. The tube was vigorously shaken and centrifuged at 1500 rpm for 5 minutes at 4°C to detach the cells from the filter. The filter was carefully removed without disturbing the pellet, followed by an additional centrifugation step for 2 minutes to firm the pellet. The Sørensen buffer was replaced with 1 ml of clean Sørensen buffer and the tube was centrifuged again for 5 min. Finally, the Sørensen buffer was discarded, and 170 μl of ice-cold RLN buffer was added.

RNA extraction was carried out using the RNeasy® Plant Mini Kit (Qiagen GmbH, Hilden, Germany) following the manufacturer’s instructions starting from step 2 and using only 300 μl ethanol in step 4. RNA concentrations were quantified using the Qubit RNA High Sensitivity Assay Kit (Thermo Fisher Scientific Inc, Germany) and Qubit 4 Fluorometer (Thermo Fisher Scientific Inc, Germany). Sequencing was performed on an Illumina NovaSeq instrument (Illumina Inc., San Diego, CA, USA) at the Cologne Center for Genomics (Köln, Germany) with 2 × 100 bp paired-end reads, polyA selection, and a sequencing depth of about 50 Mio sequences (see Supplementary Table 1).

For single-cell RNA extraction of Ebriida sp., we followed the protocol of Hagemann-Jensen et al. (2020). Single cells of Ebriida sp. were isolated from samples originating from North Slope, Alaska (71.404558 N, 156.530021 W) and grown in F4 medium with nitschoid diatoms as prey. Before RNA extraction, cells were starved for 24 hours and given in the lysis buffer with a micromanipulator. Three replicates were pooled and sequenced with an Illumina NextSeq sequencer (Illumina Inc., San Diego, CA, USA) at the Cologne Center for Genomics (Köln, Germany) with 2 × 100 bp paired-end reads, polyA selection, and a sequencing depth of about 50 Mio sequences as well (see Supplementary Table 1).

### 2.2. Transcriptome assembly

The quality of the 15 newly sequenced transcriptomes (*Rhogostoma*, *Katarium polorum*, and Ebriida sp.) as well as of *Fisculla terrestris* (Gao et al., 2025; Solbach et al., 2021) and the two Protaspidae strains (Lax et al., 2025) was assessed using FastQC v. 0.11.9 (Andrews, 2010), followed by RNA-seq error corrections with Rcorrector v 1.0.6 (Song and Florea, 2015) and quality filtering and adapter trimming with FastP v 0.23.2 (Chen et al., 2018). Potential contaminations from prokaryotes, plants, fungi, and humans were excluded using Kraken2 v 2.1.2 (Wood et al., 2019). Ribosomal RNA reads were identified using SortMeRNA v 4.3.4 (Kopylova et al., 2012) and blasted against the PR^2^ database v. 4.14.0 (Guillou et al., 2013) using BLASTN v. 2.10.0 (Camacho et al., 2009) to confirm any *Rhogostoma*/Thecofilosea sequences. The messenger RNA reads were assembled using Trinity v 2.14.0 (Grabherr et al., 2011), and candidate coding regions were identified using TransDecoder v 5.5.0 (Haas, 2018). Subsequently, the RNA-Seq read representation of the Trinity transcripts was validated using Bowtie2 v 2.4.1 (Langmead et al., 2019; Langmead and Salzberg, 2012).

We extended our data set by including 28 rhizarian transcriptomes/genomes (Balzano et al., 2015; Burki et al., 2013; Gerbracht et al., 2022; Gomaa et al., 2021; Grant et al., 2012; Keeling et al., 2014; Lhee et al., 2021; Rodríguez-Ezpeleta et al., 2007; Schwelm et al., 2015; Sierra et al., 2016) (see Supplementary Table 2). Candidate coding regions of nucleotide assemblies were identified using TransDecoder v 5.5.0 (Haas, 2018). The completeness of all assembled transcriptomes was evaluated with benchmarking universal single-copy orthologs (BUSCO) v 5.2.2 (Manni et al., 2021a, 2021b) and the eukaryote odb10 database.

### 2.3. Phylogenomic analyses

PhyloFisher v. 1.2.13 (Tice et al., 2021) was employed to identify orthologs for the muti-gene phylogeny of the Rhizaria, based on the provided database comprising 240 genes from 304 taxa across the eukaryotic tree of life. To screen for paralogs and contaminants, single gene trees of all 2490 genes were revised manually and using a customized R script. The cleaned data sets were filtered for Rhizaria transcriptomes, considering only taxa with more than 65 % amino acid coverage across all 240 genes. Exceptions were the single-cell transcriptomes of Protaspidae sp. and Ebriida sp., which had overall low gene coverage but were still included in the phylogenomic analysis. Further, the 240 genes were filtered, keeping only genes present in more than 50% of the taxa. A concatenated alignment consisting of 222 genes spanning 40 Rhizaria strains was built using PhyloFisher v. 1.2.13 (Tice et al., 2021).

The maximum likelihood (ML) tree was constructed with IQ-Tree v. 2.1.2 (Minh et al., 2020) using the site-heterogeneous mixture model LG + C60 + F + Γ and 1000 Ultra-Fast Bootstrap replicates (UFB) as well as SH-like approximate likelihood ratio test (SH-aLRT).

### 2.4. Transcriptome annotation and comparison

OrthoFinder v. 2.5.2 (Emms and Kelly, 2019, 2015) was employed to identify phylogenetic hierarchical orthogroups across all *Rhogostoma* strains, *Fisculla terrestris*, and *Katarium polorum*. Beforehand, the redundancy of the Trinity contigs was reduced by clustering at 95% identity over at least 90% of the shorter contig length using CD-HIT v. 4.8.1 (Fu et al., 2012). A customized R script was employed to filter the TransDecoder protein sequences by the clustered Trinity contigs and to split TransDecoder protein sequences with multiple predicted coding regions. In the case of overlapping coding regions, the longer one was retained. To verify the quality of the clustered contigs, we checked their completeness using BUSCO v 5.2.2 (Manni et al., 2021a, 2021b) and the eukaryotic odb10 database, and the RNA-Seq read mapping using Bowtie2 v 2.4.1 (Langmead et al., 2019; Langmead and Salzberg, 2012). Further, the gene expression was quantified with Salmon v 1.9.0 (Patro et al., 2017) in alignment-based mode. The protein sequences were functionally annotated based on the Kyoto Encyclopedia of Genes and Genomes (KEGG) database (Kanehisa, 2019; Kanehisa et al., 2023; Kanehisa and Goto, 2000) using eggNOG-mapper v. 2.1.9 (Cantalapiedra et al., 2021).

The number of shared and unique orthogroups across all strains was visualized using ggupset v. 0.4.0 (Ahlmann-Eltze, 2024). Only the top 25 combinations with the highest number of orthogroups were considered. In addition, a general overview of the relative proportions of functional annotations from 26 categories was provided by bar charts visualized with ggplot2 (Wickham, 2011). The relative proportions of functional annotations were calculated for all species, the core set of orthogroups (i.e., orthogroups shared by all strains), the two *Rhogostoma* clusters, *Katarium polorum* and *Fisculla terrestris*, respectively.

A detailed overview of the metabolic pathways of the amino acid, nucleotide, carbohydrate, and lipid metabolism was created based on the KEGG module. For this, the presence or absence of each KEGG orthology (KO) term was assessed per KEGG module, the minimum KO terms required for each module were determined, and the percentage of KO terms for each reaction step (the number of preset KO terms divided by the minimum number of KO terms required for the respective reaction) was calculated. Only KEGG modules for which (for at least one strain) 50% or more of the KO terms were present and no more than a total of three KO terms were missing were considered for the calculation of the heatmaps and the summarisation of the amino acid, central carbohydrate, and lipid metabolism modules into a graph. The graph was visualized using Cytoscape v. 3.9.0 (Shannon et al., 2003).

For the gene enrichment analysis of the two phylogenetically distinct *Rhogostoma* clusters multiple steps were conducted: First, the quantified gene expressions from salmon were summarized for each strain and orthogroup using tximport v. 1.32.0 (Soneson et al., 2016), generating matrices containing the weighted mean of the contig length, the effective contig length, the number of reads (counts) and the transcript per million (TPM) for each orthogroup. Second, low count orthogroups were excluded, keeping only orthogroups with counts per million > 1 in six or more strains. Third, principal component analysis (PCA) based on the filtered expression level of the orthogroup was calculated using the functions vst and plotPCA (DESeq2 v. 1.44.0; (Love et al., 2014)). Fourth, differential expression analysis was conducted with DESeq2 v. 1.44.0 (Love et al., 2014) comparing the two *Rhogostoma* clusters with six strains each. Orthogroups with |log2 fold change| ≥ 1 and adjusted p-value < 0.001 were considered as differentially expressed. The enrichment analysis of KO terms was carried out using GOseq v. 1.56.0 (Young et al., 2010). The enrichment analysis was performed for all 12 Rhogostoma strains, using the length and the functional assignment of the orthogroups of each strain, respectively. Only KO terms that were significantly enriched in the enrichment analysis of all strains within one Rhogostoma cluster were retained.

## 3. Results

### 3.1. Phylogenomic analysis

To shed light on the evolutionary relationships of Thecofilosea, we conducted a phylogenetic analysis based on a comprehensive multi-gene data set of 222 concatenated genes (71,479 amino acids) derived from, in total, 40 Rhizaria transcriptomes (Fig. 1, Supplementary Table 2, Supplementary Fig. 1 & 2). Among these transcriptomes are a total of 15 newly added Thecofilosea transcriptomes: 14 derived from monoclonal cultures with high gene and site coverage - 12 *Rhogostoma* (Cryomonadida) and two Tectofilosida transcriptomes, as well as one single-cell transcriptome of Ebriida sp. with moderate gene and site coverage, as it is typical for single-cell transcriptomes (Fig. 1, Supplementary Table 1).

**Figure 1:**
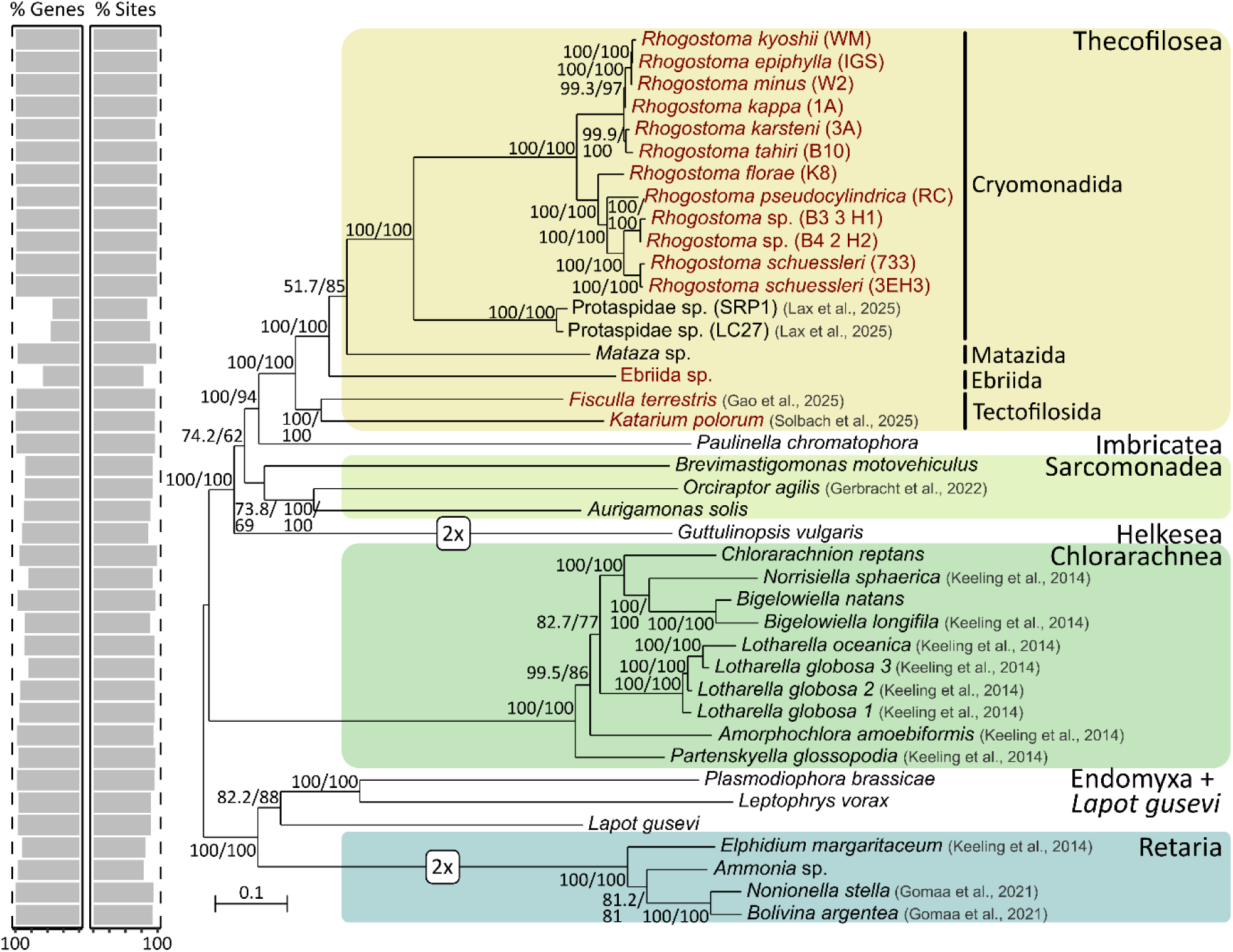
Multi-gene phylogeny of the Rhizaria. Maximum likelihood tree (LG+C60+F+G model) based on an alignment comprising 222 concatenated genes (71,479 amino acids) derived from 40 Rhizaria transcriptomes. Support values were obtained from Shimodaira–Hasegawa approximate likelihood ratio test (SH-aLRT) and 1000 ultrafast bootstraps (UFB). The percentage of gene coverage and the percentage of the site coverage of the present genes for each taxon are shown on the left. Newly added transcriptomes are highlighted in red.

The phylogenetic analyses provided a highly supported backbone of the Thecofilosea, Imbricatea, Sarcomonadea and Helkesea (Fig. 1). At the base of the Thecofilosea, *Fisculla terrestris* (Fiscullidae) and *Katarium polorum* (Chlamydophryidae) form a fully supported monophyletic group, the Tectofilosida. The Ebriida sp. branch with high support basal to Matazida. Within Cryomonadida, two distinct, fully supported monophyletic subclades were identified. The first subclade included all Protaspidae strains and clustered at the base of the second subclade, including all *Rhogostoma* strains. The *Rhogostoma* strains were further divided into two fully supported clusters, each comprising six strains: Cluster 1, characterized by short branches, included *R. kyoshii*, *R. epiphylla*, *R. minus*, *R. kappa* and *R. tahiri*, and cluster 2, with longer branches, included *R. florea*, *R. pseudocylindrica* and two strains each of *R. schuessleri* and unidentified *Rhogostoma* species.

### 3.2. Comparative transcriptomics

To explore the physiological traits of free-living the heterotrophic Thecofilosea, we compared and functionally annotated the whole transcriptomes of Thecofilosea species based on the KEGG database. Only the culture-based and thus deeply sequenced transcriptomes of *Rhogostoma*, *Fisculla*, and *Katarium* were considered, not the single-cell transcriptome of Ebriida sp. due to the comparably lower coverage of single-cell transcriptomes (Supplementary Table 1, Supplementary Figure 3). The transcriptomes comprised an average of 67,766 clustered contigs, with a mean mapping rate of 95% when aligning the reads back to the assembled contigs and a mean completeness of 86% according to benchmarking universal single-copy orthologs (BUSCO) of the Eukaryota dataset (Fig. 2B). Further, these transcriptomes contained an average of 44,972 predicted open reading frames (ORFs, Supplementary Table 1).

**Figure 2:**
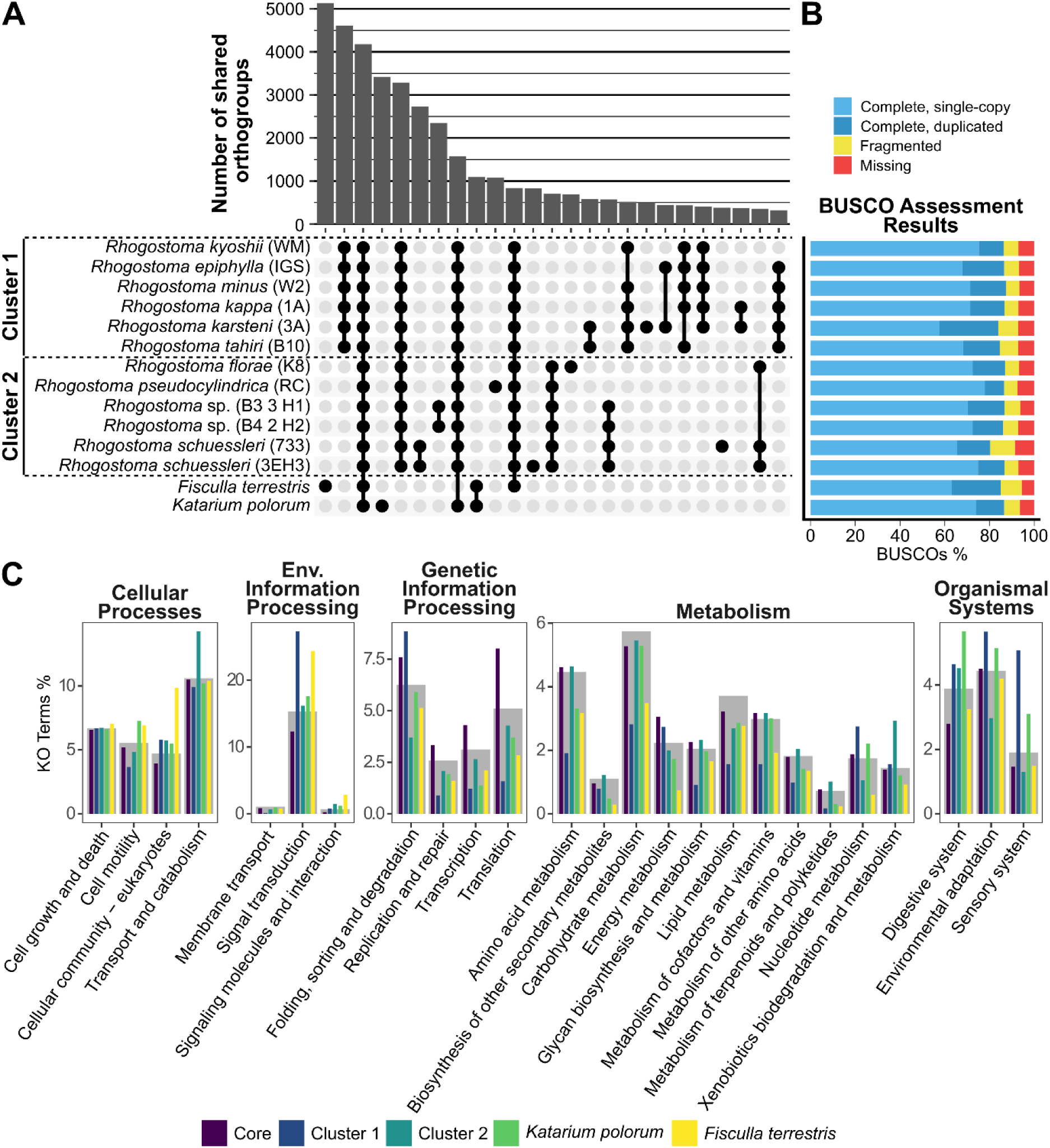
**Shared orthogroups and functional annotations of the Thecofilosea**. **(A)** The upper bar chart shows the number of shared orthogroups that are unique to the respective combination of Thecofilosida strains displayed in the matrix below. Only the top 25 strain combinations, based on the number of shared orthogroups, are shown. **(B)** BUSCOs assessment of the assembled Thecofilosida transcriptome, based on the Eukaryota database. **(C)** A selection of the functional annotations (KEGG orthology) of the Thecofilosida transcriptomes The grey bar charts in the background display the relative proportions of functional annotations of all transcriptomes for the respective category. In addition, the relative proportions of functional annotations of the core set of orthogroups (purple), the *Rhogostoma* clusters 1 (blue) and 2 (dark green), *Katarium polorum* (light green) and *Fisculla terrestris* (yellow) are shown for the respective category.

Overall, 93% of the ORFs were assigned to 58,676 orthogroups (Fig. 2A). We identified a core set of 4,178 orthogroups that was shared by all investigated Thecofilosea species (Fig. 2A). Compared to the functional annotations of all orthogroups, the core orthogroups exhibited a higher relative proportion of KO terms related to genetic information processing (Fig. 2C), in particular, related to replication and repair, transcription and translation. The two phylogenetically distinct *Rhogostoma* clusters showed equally distinct sets of orthogroups. Cluster 1 shared a higher number of unique orthogroups (4,612) than cluster 2 (704, Fig. 2A). Further, cluster 1 comprised a higher proportion of KO terms related to signal transduction, sensory systems, and environmental adaptation. In contrast, Cluster 2 comprised a higher proportion of KO terms related to transport and catabolism compared to the overall functional annotation (Fig. 2C). Notably, both Tectofilosida species, *Fisculla terrestris* and *Katarium polorum* exhibited high numbers of unique orthogroups, 5,134 and 3,417, respectively (Fig. 2A). Both species showed a higher relative proportion of KO terms involved in cell motility. *Fisculla terrestris* additionally showed a relative increase of KO terms associated with cellular community processes such as focal adhesion, adherens junctions, tight junctions, and gap junctions compared to the overall functional annotation (Fig. 2C).

To gain insights into the metabolic adaptations of these free-living, heterotrophic Thecofilosea strains, we analyzed KEGG modules involved in amino acid, carbohydrate, nucleotide, and lipid metabolism (Fig. 3) and reconstructed the pathways for central carbohydrate and amino acid metabolism (Fig. 4 & 5, Supplementary Figures 4-29). The majority of genes involved in the central carbohydrate metabolism were expressed, although the complete pyruvate oxidation pathway could only be reconstructed for 6 out of 14 strains. The *Rhogostoma* cluster 2 lacked few genes for the non-oxidative pentose phosphate pathway. Furthermore, most of the genes involved in nucleotide biosynthesis were present throughout the dataset, except for genes related to the *de novo* purine biosynthesis, which were exclusively found in *Katarium polorum*. In contrast to the relatively conserved central carbohydrate and nucleotide metabolisms, greater variability was observed in amino acid and lipid metabolism. All strains expressed most genes involved in the biosynthesis of proline, valine, isoleucine, cysteine, methionine, serine, threonine, glutamate, and glycine. However, only *Fisculla terrestris* and *Katarium polorum* appeared to synthesize arginine, tryptophan, histidine, and lysine. *Fisculla terrestris* additionally possessed most genes to synthesize leucine, phenylalanine, and tyrosine. Differences in the expression of genes involved in lipid metabolism between the strains were detected, particularly genes involved in sterol biosynthesis. For example, strains belonging to the *Rhogostoma* Cluster 2 expressed a greater number of enzymes involved in the biosynthesis of C18/19/21-steroids.

**Figure 3:**
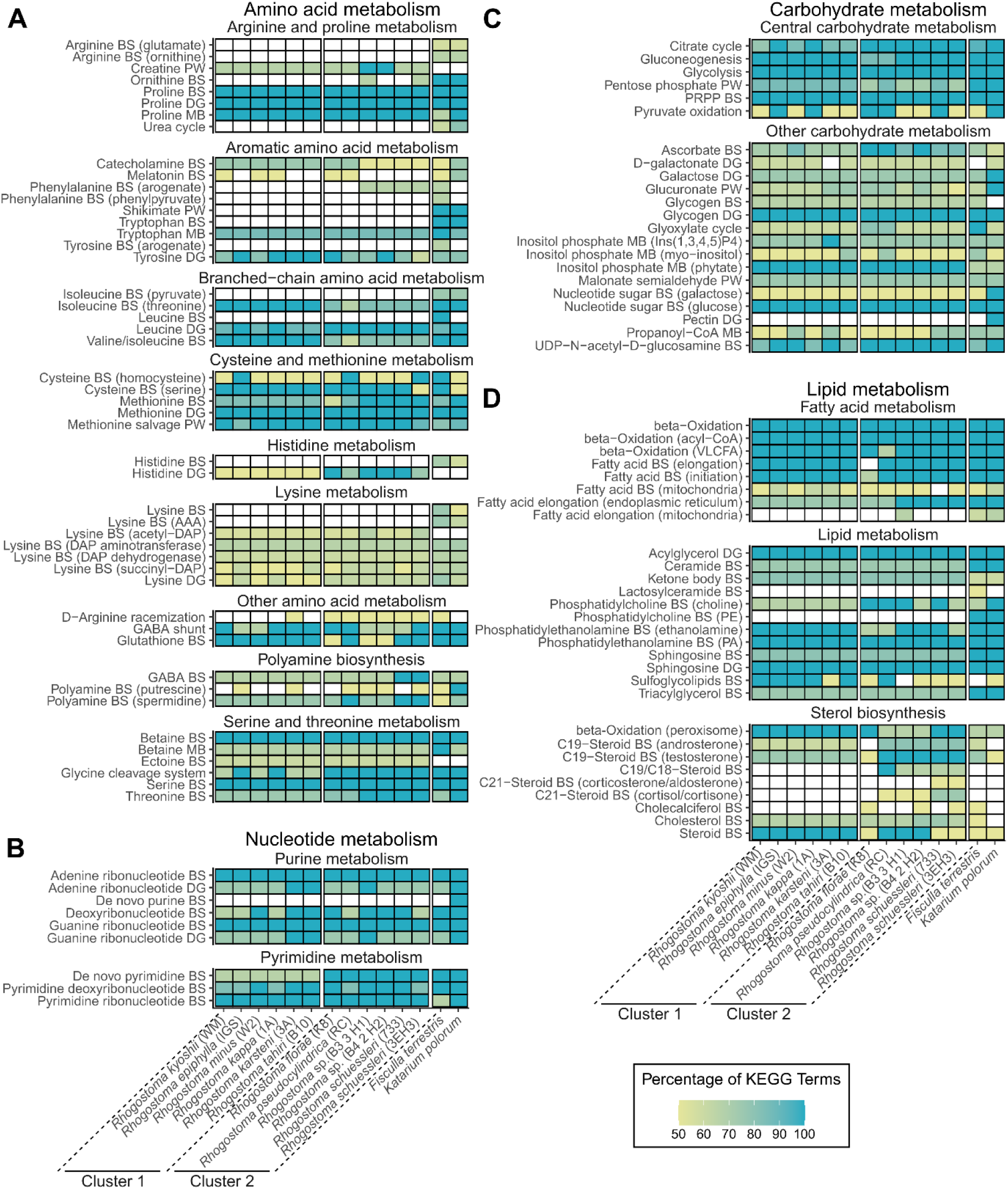
Overview of metabolic pathways for the Thecofilosea transcriptomes based on KEGG modules. The heat maps show the presence and absence of the KEGG modules for amino acid metabolism (A), nucleotide metabolism (B), carbohydrate metabolism (C) and lipid metabolism (D) for each culture-based Thecofilosea transcriptome. The colour indicates the percentage of KO terms present, normalised to the minimum number of KO terms required for each module. White boxes indicate that either more than 50% or more than three of the required KO terms were missing. Modules that were absent in all Thecofilosea transcriptomes are not shown. Abbreviations: BS Biosynthesis, PW Pathway, MB Metabolism, DG Degradation.

**Figure 4:**
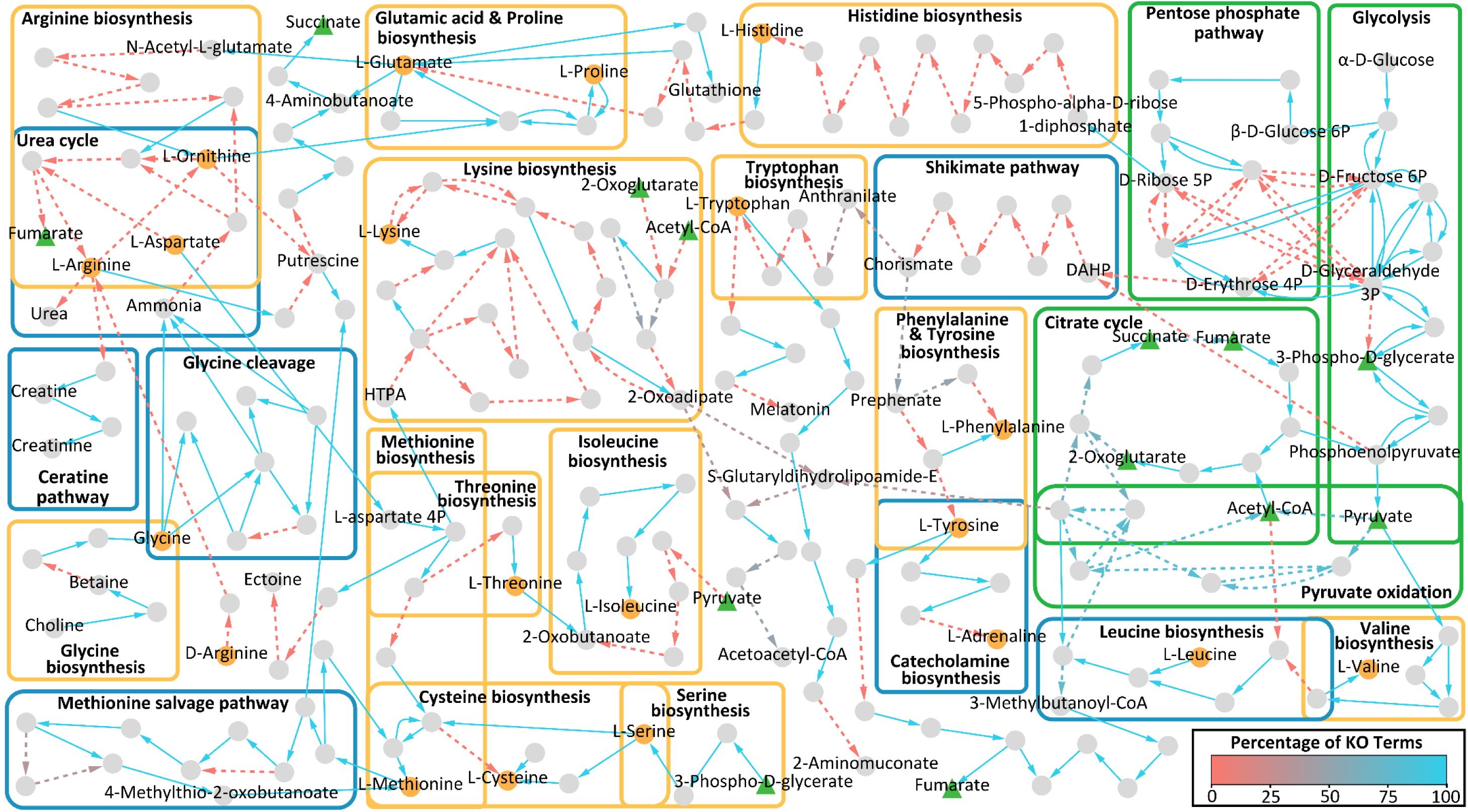
Overview of the central carbohydrate and amino acid metabolism for *Rhogostoma kyoshii* (WM). The graph illustrates a reconstruction of the central carbohydrate (green boxes) and amino acid metabolism for *Rhogostoma kyoshii* (WM) based on KEGG ontologies and KEGG modules. Nodes represent components and edges represent enzymatic reactions. Amino acids are highlighted in orange, central compounds of carbohydrate metabolism in green. The edge color indicates the percentage of KO terms present, normalized to the minimum number of KO terms required for the respective reaction. Solid edges indicate that all KO terms were present for the respective reaction.

### 3.3. Enrichment analysis

In addition to the analyses of presence-absence data, we compared the relative expression patterns of the two *Rhogostoma* clusters based on 20,000 orthogroups. A PCA showed a clear differentiation of *Rhogostoma* clusters 1 and 2, explaining 82% of the total variation (Fig. 6A). In addition, differences in the expression patterns of the long-branched, i.e., evolutionary distant *Rhogostoma* cluster 2 (Fig. 1) explained 5% of the total variation, with strains of the same species grouping together.

**Figure 5:**
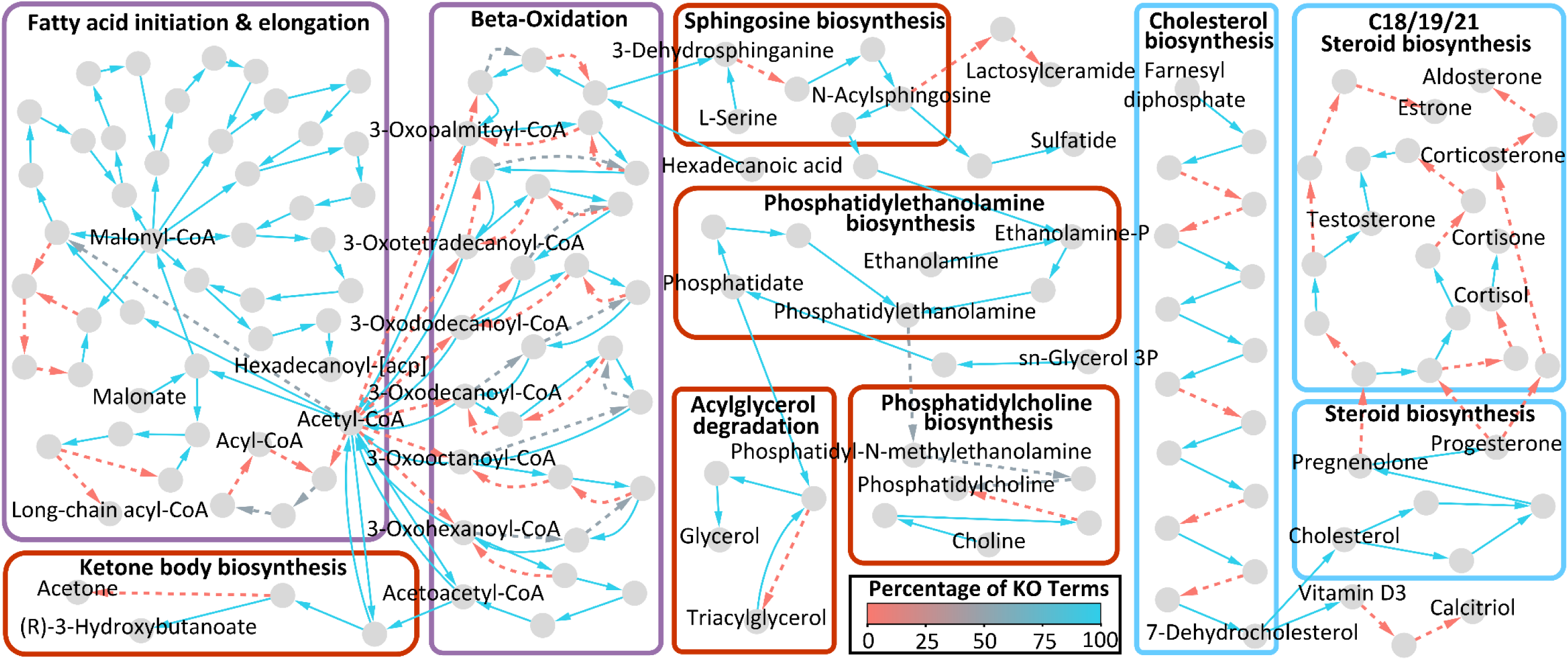
Overview of the lipid metabolism for *Rhogostoma kyoshii* (WM). The graph illustrates a reconstruction of the fatty acid (purple boxes), sterol (blue boxes), and lipid (red boxes) metabolism for *Rhogostoma kyoshii* (WM) based on KEGG ontologies and KEGG modules. Nodes represent components and edges represent enzymatic reactions. The edge color indicates the percentage of KO terms present, normalized to the minimum number of KO terms required for the respective reaction. Solid edges indicate that all KO terms were present for the respective reaction.

**Figure 6:**
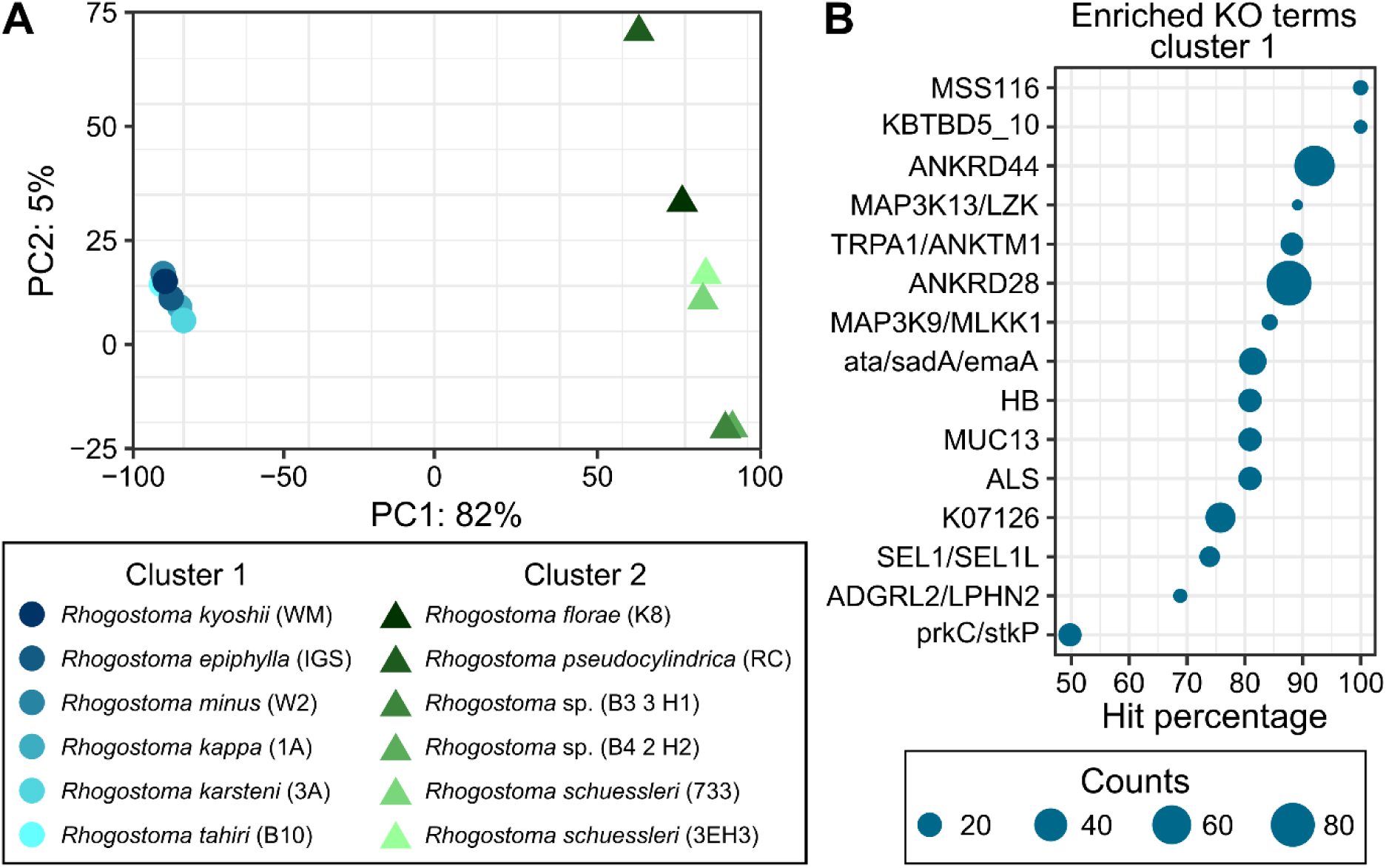
Comparative transcriptomics of *Rhogostoma*. **(A)** Principal component analysis of all *Rhogostoma* strains based on the expression patterns of 20,000 orthogroups. The color and shape of the points encode for *Rhogostoma* clusters 1 (blue shades, round) and 2 (green shades, triangular). **(B)** Significantly enriched KO terms in all six *Rhogostoma* strains of cluster 1. The enrichment analysis was based on 7,502 out of 20,000 differentially expressed orthogroups (adjusted p-value < 0.001, |log2 fold change| ≥ 1). The size of the points indicates the frequency of the respective KO terms in the higher expressed orthogroups of *Rhogostoma* cluster 1. The hit percentage describes the ratio of KO terms in higher expressed orthogroups of *Rhogostoma* cluster 1 compared to all orthogroups.

Differential expression analysis revealed 7,502 out of the 20,000 orthogroups to be differentially expressed. About one-third of these orthogroups could be functionally annotated. A subsequent enrichment analysis identified a total of 15 KO terms that were consistently enriched across all six species of *Rhogostoma* cluster 1 (Fig. 6B). In contrast, no KO terms were consistently enriched across all six species of *Rhogostoma* cluster 2. ANKRD28 and ANKRD44 (Ankyrin repeat domains 28 and 44) were the most frequent KO terms of the enriched KO terms of *Rhogostoma* cluster 1, with 84 and 66 counts, respectively. ANKRD28 and ANKRD44 occurred almost exclusively (∼90%) in the significantly enriched orthogroups (hit percentage, Fig. 6B). Furthermore, ata/sadA/emaA (trimeric autotransporter adhesin), ALS (agglutinin-like protein), MUC13 (mucin 13), and TRPA1/ANKTM1 (transient receptor potential cation channel subfamily A member 1) were identified among the enriched KO terms of *Rhogostoma* cluster 1. They also exhibited a high prevalence, with 26, 18, 18, and 17 counts, respectively, and ∼80% hit percentage in the significantly enriched orthogroups (Fig. 6B).

## 4. Discussion

### 4.1. The physiological capacity of Thecofilosea (Cecozoa)

We identified a core set of conserved genes and pathways that were shared across all investigated Thecofilosea species. First, we identified a core set of shared orthogroups that included an exceptionally high proportion of genes associated with genetic information processing, i.e., translation, transcription, and replication. As these genes are essential for all living cells and are known to be highly conserved (Yao and O’Donnell, 2016), it was expected that these genes would be overrepresented in the core set of orthogroups. Second, we show an overall high completeness in the central carbohydrate and nucleotide metabolism, including the presence of glycolysis, the citrate cycle, and the capacity to synthesize all nucleotides. The few missing enzymes can most likely be explained by differences in the homologs due to rapid evolutionary rates or incompleteness of the transcriptome data. Overall, the robust identification of conserved genes and pathways emphasizes the soundness of our methods and data.

We were able to reconstruct the *de novo* synthesis for at least nine amino acids for all investigated Thecofilosea species. However, we also found variations in the physiological traits of the species. Heterotrophic microorganisms usually lack pathways for synthesizing certain amino acids and consequently depend on salvaging them from their prey. For example, the predatory amoebae *Dictyostelium* and *Arcella* have lost the capability to synthesize 11 and 5 amino acids, respectively (Payne and Loomis, 2006; Ribeiro et al., 2020). Our data indicate the absence of numerous amino acid pathways, for instance, tryptophan and histidine, particularly in *Rhogostoma* – indicating their dependence on the uptake of these compounds from their prey.

### 4.2. Physiology in an evolutionary context

4.2.1. **Multi-gene phylogeny**

By incorporating the new Thecofilosea transcriptomes into existing public Rhizaria data, our phylogenetic analysis expands the latest Cercozoa multi-gene phylogeny of Irwin et al. (2019). In our phylogenetic tree, the superclass Ventrifilosa is highly supported – a hypothesized monophylum of the predominantly shell-bearing and free-living cecozoan groups Thecofilosea and Imbricatea (Cavalier-Smith and Karpov, 2012). We further confirm that the Tectofilosida are indeed monophyletic as suggested by Dumack et al. (2017). Lastly, the Sarcomonadea branch basal to Ventrifilosa, as indicated by numerous single-gene phylogenies (Howe et al., 2011; Scoble and Cavalier-Smith, 2014).

#### 4.2.2. Orthogroup similarity reflects the phylogenetic distance

The high coverage and saturation of our culture-derived transcriptomic data allowed for whole transcriptome comparisons of physiological traits among Thecofilosea. The physiological capabilities based on nearly complete transcriptomes highly reflect evolutionary distance across all studied Thecofilosea strains. Within Tectofilosida, although only represented by two strains, the high evolutionary distance – indicated by long branches in the phylogenetic analyses – was reflected by the high number of unique orthogroups in each species and a moderate amount of shared orthogroups. Notably, *Fisculla terrestris* exhibited a higher proportion of unique orthogroups related to cell adherence, fusion, and cell-to-cell communication, likely reflecting its nature of frequent fusion (Gao et al., 2025).

*Rhogostoma* radiated into two clusters with notable differences in both orthogroups and gene expression patterns. The short-branched, i.e., evolutionary close, *Rhogostoma* cluster 1 exhibited a high number of shared orthogroups and clustered closer together in the PCA, compared to the long-branched, i.e., evolutionary distant, *Rhogostoma* cluster 2. Remarkably, *Rhogostoma* cluster 1 showed a higher proportion of genes related to signal transduction, sensory systems, and environmental adaptation in the orthogroups unique to this cluster. In addition, differences in gene expression patterns revealed enrichment in genes in *Rhogostoma* cluster 1 associated with first, sensory processes, including the reception of heat, pain, or environmental irritants (TRPA1/ANKTM1, Bautista et al., 2006), second protection and lubrication of cell surfaces (Muc13, Williams et al., 2001), third, cell migration and focal adhesion formation (ANKRD28, Tachibana et al., 2009), and fourth, cell adhesion to biotic and abiotic surfaces and biofilm formation (ata/sadA/emaA and ALS, Bentancor et al., 2012; Mintz, 2004; Oh et al., 2019; Raghunathan et al., 2011).

Aside from inter-cluster variation, there is additionally a large inter-specific variation in *Rhogostoma*, showcasing that even closely related and morphologically similar species exhibit distinct physiological traits and thus, potentially distinct ecological influence. It is important to note that although transcriptomic responses exhibit a snapshot of metabolic activity (Martin and Wang, 2011; Raghavan et al., 2022), we analyzed a high density of input cells reflecting the transcriptomic response of thousands of individuals, minimizing temporal and individual variation. To further increase the comparability, we grew all *Rhogostoma* strains in the same medium, strengthening our interpretation of the results.

#### 4.2.3. Reduced gene set of *Rhogostoma*

In addition to a core set of orthogroups, each investigated evolutionary clade expressed unique transcripts, providing evidence that protistan functions in ecosystems cannot be easily generalized, even at a low taxonomic level. We show that *Rhogostoma*, a genus containing morphologically highly similar species (Öztoprak et al., 2020) and being the most derived clade in our phylogenetic tree, exhibited an exceptionally high variability in its physiological repertoire. The question arises, which selective forces led to the evolution of different physiological traits in species with highly similar morphologies?

As our phylogenetic tree shows, the Rhogostomidae (Cryomonadida) are closely related to the Protaspidae (Cryomonadida), highly specialized parasites of algae (Drebes et al., 1996; Schnepf and Kühn, 2000). The transition from a free-living to a parasitic lifestyle usually leads to the loss of existing functions and a simplified metabolism, as parasites exploit the resources of their host (Jackson et al., 2016; Poulin and Randhawa, 2015). Thus, the transition to parasitism is often thought to be irreversible, yet several studies, for example on diplomonads, provide evidence for a secondary free-living lifestyle (Wiśniewska et al., 2024; Xu et al., 2016). We hypothesize that the ancestors of the Cryomonadida underwent a loss of functional and genomic diversity with the specialization towards parasitism and that the ancestor of the Rhogostomidae broadened the prior narrow prey spectrum and specialized on feeding on bacteria in addition to eukaryotes.

Horizontal gene transfer (HGT) is, in general, considered to be a minor driver of eukaryote evolution (Keeling, 2024, 2019). Instead, eukaryotes are considered mainly to evolve by gene duplication and the subsequent adaption of homologs to different functions. Nonetheless, there is evidence that HGT in microbial eukaryotes may represent an escape from increasing adaptation to parasitism, which typically involves a loss of genomic and functional diversity (Wiśniewska et al., 2024; Xu et al., 2016), i.e., that HGT may contribute to the reversibility of parasitism and readaptation to a free-living lifestyle through the acquisition of new functions.

Overall, the potential back-transition of *Rhogostoma* from parasitism to free-living might have caused the acquisition of even single functional genes to lead to a new speciation event, thus contributing to the current remarkable species diversity of *Rhogostoma*.

## 5. Conclusion

The outstanding high coverage of the transcriptomes and the deeply sampled phylogeny of the free-living and heterotrophic Thecofilosea allowed us to draw conclusions on their physiological capabilities and compare them in an evolutionary context. The Thecofilosea possess a core set of conserved genes involved in genetic information processing, nucleotide metabolism, and central carbohydrate metabolism. However, the high variability in amino acid and lipid metabolism, as well as the differentially expressed and enriched genes, indicate potentially distinct functional roles. These findings emphasize the remarkable physiological diversity even among closely related, morphologically highly similar taxa. Considering that the Thecofilosea represent only a fraction of the tremendous protistan diversity, the need for further research is evident. Our methodological approach paves the way for subsequent studies on a larger scale.

## 7. Data Accessibility

The raw sequencing data are available under NCBI BioProject PRJNA1262159 and the trinity transcriptomes will be available on Zenodo (DOI: https://doi.org/10.5281/zenodo.15091355) upon publication.

## 8. Funding

This study was supported by grants from the German Research Foundation (DFG) in the framework of the priority program SPP 1991: TAXON-OMICS (Project IDs 447013012 and 221301018) and with the grant number 555596351.

## Acknowledgements

We would like to thank Lukas Isenhart for carrying out most of the RNA extractions and Gordon Lax for providing early access to the single-cell sequences of Protaspida. We furthermore thank the Regional Computing Center of the University of Cologne (RRZK) for providing computing time on the DFG-funded (Funding number: INST 216/512/1FUGG) High Performance Computing (HPC) system CHEOPS as well as support.

## 10. Authors’ contributions

KD, MS and MB designed the study. NJME established and performed the protocol for single-cell RNA extraction of the Ebriida strain. JF created all figures and performed the bioinformatical and statistical analyses. JF outlined the manuscript, all co-authors commented on the manuscript and approved the submitted version.

## 11. Competing interests

The authors declare no competing interests.

## Notes

### Competing Interest Statement

The authors have declared no competing interest.

