## Supplementary data for "Physiological diversity and adaptation of Rhizaria revealed by phylogenomics and comparative transcriptomics"

**Supplementary Table 1: Overview of transcriptome assemblies.** The table shows the number of raw, trimmed and filtered reads of all *Rhogostma* strains, *Fisculla terrestris*, *Katarium polorum*, Protaspida sp. and Ebriida sp. In addition, the number of Trinity contigs and TransDecoder peptides is given, as well as the BUSCO completeness (Eukaryote odb10 database) and the Bowtie RNA-Seq read mapping rate. The latter is given for both the original and clustered Trinity contigs for all *Rhogostma* strains, *Fisculla terrestris* and *Katarium polorum*.

| Species | Raw reads | Reads after trimming | Reads after contamination filtering | mRNA reads | Clustered/Not Clustered | Trinity | | | BUSCO | | Mapping rate | TransDecoder | | |
|---|---|---|---|---|---|---|---|---|---|---|---|---|---|---|
| | | | | | | Transcripts | N50 | Median length | Complete genes | Fragmented genes | | Peptides | N50 | Median length |
| *Rhogostoma kyoshii* (WM) | 66533991 | 53579788 | 51070053 | 50742478 | Not Clustered | 70500 | 1700 | 749 | 86.2 | 6.7 | 99.64 | 54146 | 501 | 251 |
| | | | | | Clustered | 59087 | 1661 | 725 | 86.3 | 6.7 | 95.94 | 43860 | 488 | 247 |
| *Rhogostoma epiphylla* (IGS) | 65487075 | 52520473 | 49620845 | 49201702 | Not Clustered | 87591 | 1683 | 692 | 87.9 | 6.3 | 99.63 | 65641 | 504 | 243 |
| | | | | | Clustered | 71356 | 1613 | 655 | 86.6 | 6.7 | 95.68 | 51349 | 475 | 232 |
| *Rhogostoma minus* (W2) | 66314131 | 58020320 | 55233041 | 54923000 | Not Clustered | 77982 | 1746 | 756 | 87.5 | 5.9 | 99.62 | 60335 | 512 | 251 |
| | | | | | Clustered | 64673 | 1681 | 731 | 87.5 | 5.9 | 96.13 | 48384 | 488 | 244 |
| *Rhogostoma kappa* (1A) | 51545724 | 40019834 | 37599379 | 37340949 | Not Clustered | 78813 | 1685 | 697 | 87.5 | 5.5 | 99.5 | 58373 | 497 | 245 |
| | | | | | Clustered | 64710 | 1648 | 692 | 86.7 | 6.3 | 95.46 | 47114 | 480 | 241 |
| *Rhogostoma karsteni* (3A) | 51003791 | 35252147 | 33337297 | 32956763 | Not Clustered | 112600 | 1547 | 656 | 84.3 | 8.6 | 99.29 | 81278 | 434 | 227 |
| | | | | | Clustered | 91974 | 1515 | 638 | 83.9 | 9 | 92.01 | 64913 | 414 | 220 |
| *Rhogostoma tahiri* (B10) | 51142494 | 36827766 | 34794910 | 34503204 | Not Clustered | 78688 | 1608 | 724 | 85.1 | 7.8 | 99.56 | 59521 | 467 | 241 |
| | | | | | Clustered | 65703 | 1575 | 709 | 84.7 | 8.2 | 93.72 | 48624 | 452 | 236 |
| *Rhogostoma florae* (K8) | 62748138 | 50695419 | 48711042 | 48375794 | Not Clustered | 58503 | 1876 | 784 | 87.4 | 5.9 | 99.33 | 40058 | 605 | 312 |
| | | | | | Clustered | 49440 | 1817 | 724 | 86.6 | 5.9 | 96.18 | 32067 | 590 | 307 |
| *Rhogostoma pseudocylindrica* (RC) | 71362011 | 55869020 | 54019464 | 53442745 | Not Clustered | 70970 | 1808 | 721 | 89.1 | 5.1 | 99.4 | 48490 | 565 | 285 |
| | | | | | Clustered | 56109 | 1764 | 729 | 87 | 6.3 | 96.58 | 37473 | 549 | 286 |
| *Rhogostoma* sp. (B3 3 H1) | 57476393 | 46776628 | 45348061 | 44878690 | Not Clustered | 77985 | 1910 | 898 | 87.8 | 5.9 | 99.38 | 54185 | 573 | 296 |
| | | | | | Clustered | 65178 | 1858 | 822 | 86.7 | 7.1 | 95.2 | 42393 | 551 | 285 |
| *Rhogostoma* sp. (B4 2 H2) | 52974991 | 41960462 | 40494773 | 40170318 | Not Clustered | 65232 | 1819 | 881 | 87.8 | 5.5 | 99.51 | 45318 | 572 | 302 |
| | | | | | Clustered | 54605 | 1769 | 804 | 86.2 | 6.7 | 95.13 | 35310 | 556 | 293 |
| *Rhogostoma schuessleri* (733) | 48558834 | 36144031 | 34838367 | 34125325 | Not Clustered | 74686 | 1471 | 706 | 81.6 | 9.8 | 98.53 | 52855 | 451 | 258 |
| | | | | | Clustered | 61976 | 1441 | 637 | 80.4 | 11 | 93.1 | 41357 | 444 | 250 |
| *Rhogostoma schuessleri* (3EH3) | 67459726 | 53092251 | 50988304 | 50202712 | Not Clustered | 91664 | 1760 | 771 | 87.9 | 5.1 | 98.8 | 62174 | 548 | 285 |
| | | | | | Clustered | 71423 | 1619 | 677 | 86.7 | 6.3 | 94.38 | 44303 | 503 | 271 |
| *Fisculla terrestris* | 60284486 | 54248087.7 | 50390734.7 | 50038010.8 | Not Clustered | 105351 | 1701 | 574 | 86.6 | 8.2 | 99.63 | 66945 | 534 | 262 |
| | | | | | Clustered | 91295 | 1572 | 534 | 85.1 | 9.4 | 96.64 | 54448 | 494 | 250 |
| *Katarium polorum* | 64929398 | 53808760 | 49336388 | 48736075 | Not Clustered | 92384 | 1500 | 435 | 88.2 | 5.9 | 98.72 | 46980 | 529 | 266 |
| | | | | | Clustered | 81202 | 1436 | 401 | 86.6 | 7.1 | 92.49 | 38010 | 523 | 261 |
| Ebriida sp. | 65095457 | 55622027 | 50593796 | 46007110 | Not Clustered | 32903 | 645 | 435 | 10.2 | 19.2 | 89.83 | 16924 | 221 | 171 |
| Protaspidae sp. (SRP1) | 10703734 | 7585655 | 6116460 | 5297090 | Not Clustered | 14114 | 911 | 461 | 7.4 | 5.9 | 98.35 | 6085 | 298 | 197 |
| Protaspidae sp. (LC27) | 7428223 | 5183399 | 4167035 | 3694243 | Not Clustered | 12167 | 939 | 479 | 9 | 5.9 | 97.97 | 5720 | 290 | 196 |

**Supplementary Table 2: Overview of all Rhziaria strains included in the phylogenetic analysis.** The table provides information on the data type and processing status (raw data, assembly or protein sequences), as well as the data origin, i.e., database and ID, and the citation and DOI of the corresponding publication (if available). The Rhizaria species included in the final multi-gene phylogeny are marked with an asterisk.

| Species | Type | Status | Database | Database ID | Author | DOI |
|---|---|---|---|---|---|---|
| ***Amorphochlora amoebiformis**** | Transcriptome | Assembly | Zenodo | MMETSP0042_2 | Keeling et al., 2014 | https://doi.org/10.1371/journal.pbio.1001891 |
| ***Amphilonche elongata*** | Transcriptome | Assembly | Zenodo | MMETSP1359 | Balzano et al., 2015 | https://doi.org/10.3389/fmicb.2015.00098 |
| ***Astrolonche serrata*** | Transcriptome | Proteins | EukProt v3 | EP00494 | Sierra et al., 2016 | https://doi.org/10.1093/molbev/msv340 |
| ***Aulacantha scolymantha*** | Transcriptome | Assembly | EukProt v3 | EP00463 | Balzano et al., 2015 | https://doi.org/10.3389/fmicb.2015.00098 |
| ***Bigelowiella longifila**** | Transcriptome | Assembly | Zenodo | MMETSP1359 | Keeling et al., 2014 | https://doi.org/10.1371/journal.pbio.1001898 |
| ***Bolivina argentea**** | Transcriptome | Proteins | EukProt v3 | EP01084 | Gomaa et al., 2021 | https://doi.org/10.1126/sciadv.abf1586 |
| ***Brizalina* sp.** | Transcriptome | Assembly | EukProt v3 | EP00481 | Sierra et al., 2016 | https://doi.org/10.1093/molbev/msv340 |
| ***Bulimina marginata*** | Transcriptome | Assembly | EukProt v3 | EP00487 | Sierra et al., 2016 | https://doi.org/10.1093/molbev/msv340 |
| **Protaspidae sp. (SRP1)*** | Transcriptome | RNA-Seq | NCBI | SRR31106255 | Lax et al., 2025 | https://doi.org/10.21203/rs.3.rs-7584520/v1 |
| **Protaspidae sp. (LC27)*** | Transcriptome | RNA-Seq | NCBI | SRR31106254 | Lax et al., 2025 | https://doi.org/10.21203/rs.3.rs-7584520/v1 |
| ***Elphidium margaritaceum**** | Transcriptome | Assembly | Zenodo | MMETSP1385 | Keeling et al., 2014 | https://doi.org/10.1371/journal.pbio.1001899 |
| ***Filoreta tenera*** | Transcriptome | Proteins | EukProt v3 | EP00478 | Grant et al., 2012 | https://doi.org/10.2478/prge-2012-0002 |
| ***Fisculla terrestris*** | Transcriptome | RNA-Seq | NCBI | PRJNA1108686 | Gao et al., 2025 | https://doi.org/10.1186/s12915-025-02246-3 |
| ***Globobulimina turgida*** | Transcriptome | Proteins | EukProt v3 | EP00483 | Sierra et al., 2016 | https://doi.org/10.1093/molbev/msv340 |
| ***Lotharella globosa* 1*** | Transcriptome | Assembly | Zenodo | MMETSP0041_2 | Keeling et al., 2014 | https://doi.org/10.1371/journal.pbio.1001890 |
| ***Lotharella globosa* 2*** | Transcriptome | Assembly | Zenodo | MMETSP0111_2 | Keeling et al., 2014 | https://doi.org/10.1371/journal.pbio.1001892 |
| ***Lotharella globosa* 3*** | Transcriptome | Assembly | Zenodo | MMETSP0112_2 | Keeling et al., 2014 | https://doi.org/10.1371/journal.pbio.1001893 |
| ***Lotharella oceanica**** | Transcriptome | Assembly | Zenodo | MMETSP0040_2 | Keeling et al., 2014 | https://doi.org/10.1371/journal.pbio.1001889 |
| ***Mikrocytos mackini*** | Transcriptome | Proteins | EukProt v3 | EP00477 | Burki et al., 2013 | https://doi.org/10.1016/j.cub.2013.06.033 |
| ***Minchinia chitonis*** | Transcriptome | Assembly | Zenodo | MMETSP0186 | Keeling et al., 2014 | https://doi.org/10.1371/journal.pbio.1001895 |
| ***Nonionella stella**** | Transcriptome | Proteins | EukProt v3 | EP01083 | Gomaa et al., 2021 | https://doi.org/10.1126/sciadv.abf1586 |
| ***Nonionellina* sp.** | Transcriptome | Assembly | EukProt v3 | EP00488 | Sierra et al., 2016 | https://doi.org/10.1093/molbev/msv340 |
| ***Norrisiella sphaerica**** | Transcriptome | Assembly | Zenodo | MMETSP0113_2 | Keeling et al., 2014 | https://doi.org/10.1371/journal.pbio.1001894 |
| ***Orciraptor agilis**** | Transcriptome | Assembly | ENA | HBWT01000000 | Gerbracht et al., 2022 | https://doi.org/10.1016/j.cub.2022.05.049 |
| ***Paracercomonas marina*** | EST | Proteins | EukProt v3 | EP00460 | Rodríguez-Ezpeleta et al., 2007 | https://doi.org/10.1016/j.cub.2007.07.036 |
| ***Partenskyella glossopodia**** | Transcriptome | Assembly | Zenodo | MMETSP1318 | Keeling et al., 2014 | https://doi.org/10.1371/journal.pbio.1001897 |
| ***Paulinella micropora*** | Genome | Proteins | EukProt v3 | EP00808 | Lhee et al., 2021 | https://doi.org/10.1093/molbev/msaa206 |
| ***Phyllostaurus siculus*** | Transcriptome | Proteins | EukProt v3 | EP00493 | Sierra et al., 2016 | https://doi.org/10.1093/molbev/msv340 |
| ***Rosalina sp.*** | Transcriptome | Assembly | Zenodo | MMETSP0190_2 | Keeling et al., 2014 | https://doi.org/10.1371/journal.pbio.1001896 |
| ***Spongosphaera streptacantha*** | EST | Assembly | EukProt v3 | EP00497 | Balzano et al., 2015 | https://doi.org/10.3389/fmicb.2015.00098 |
| ***Spongospora subterranea*** | Transcriptome | Proteins | EukProt v3 | EP00474 | Schwelm et al., 2015 | https://doi.org/10.1038/srep11153 |

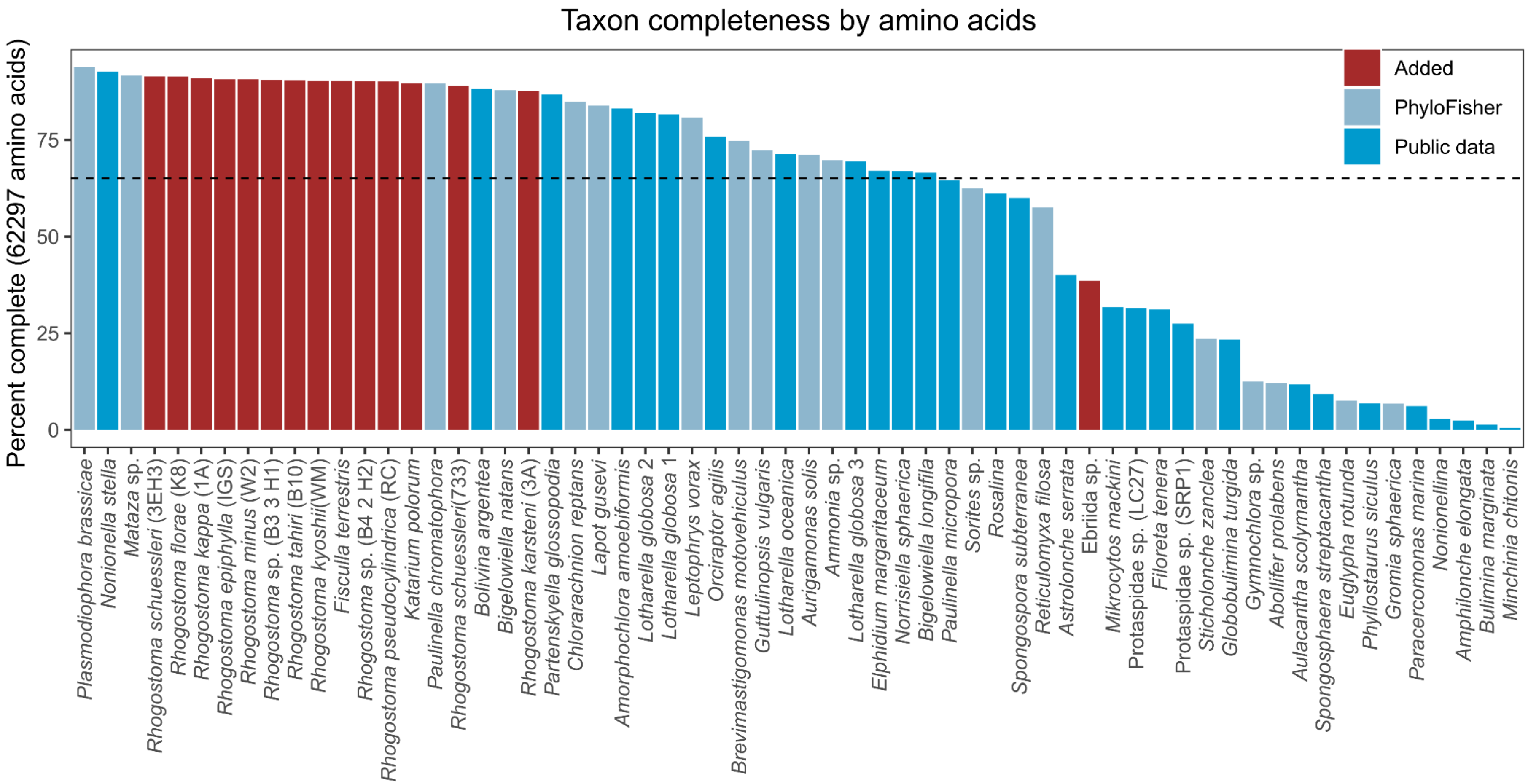


**Supplementary Figure 1: Taxon completeness by amino acids.** The bar charts show the completeness, measured as the percentage of amino acids, of a total of 240 genes selected for phylogenetic analysis, from 62 Rhizaria strains. The colours indicate the origin of the strains, i.e., newly acquired data (red), publicly available data (dark blue) and data provided by PhyloFisher (light blue).

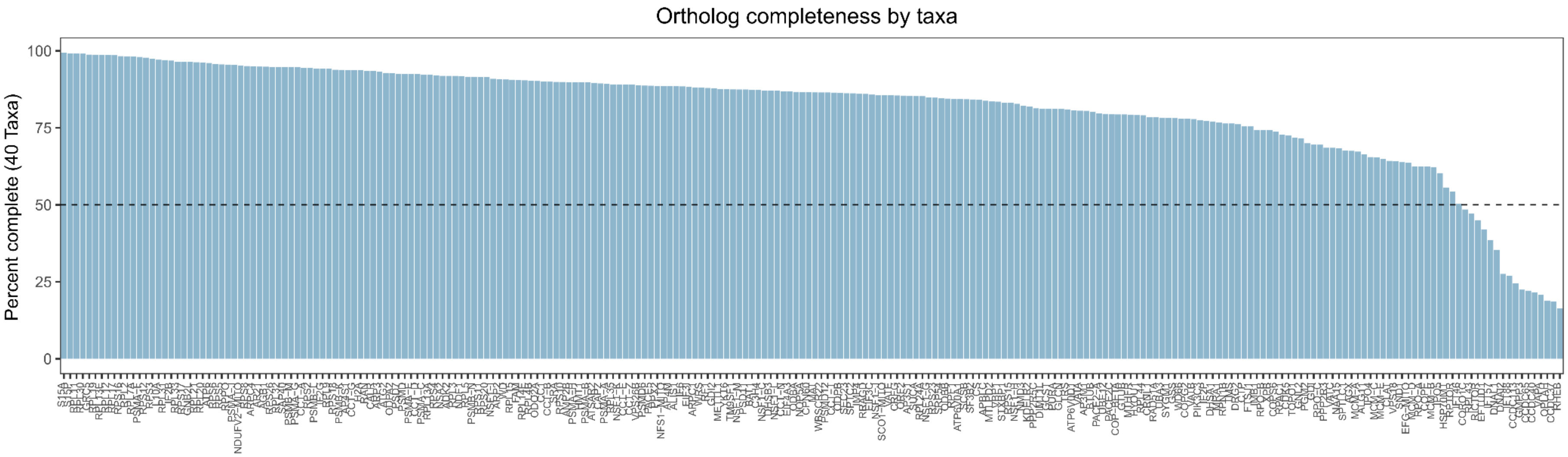


**Supplementary Figure 2: Ortholog completeness by amino acids.** The bar charts show the percentage coverage of 40 Rhizaria strains for 240 genes selected for phylogenetic analysis.

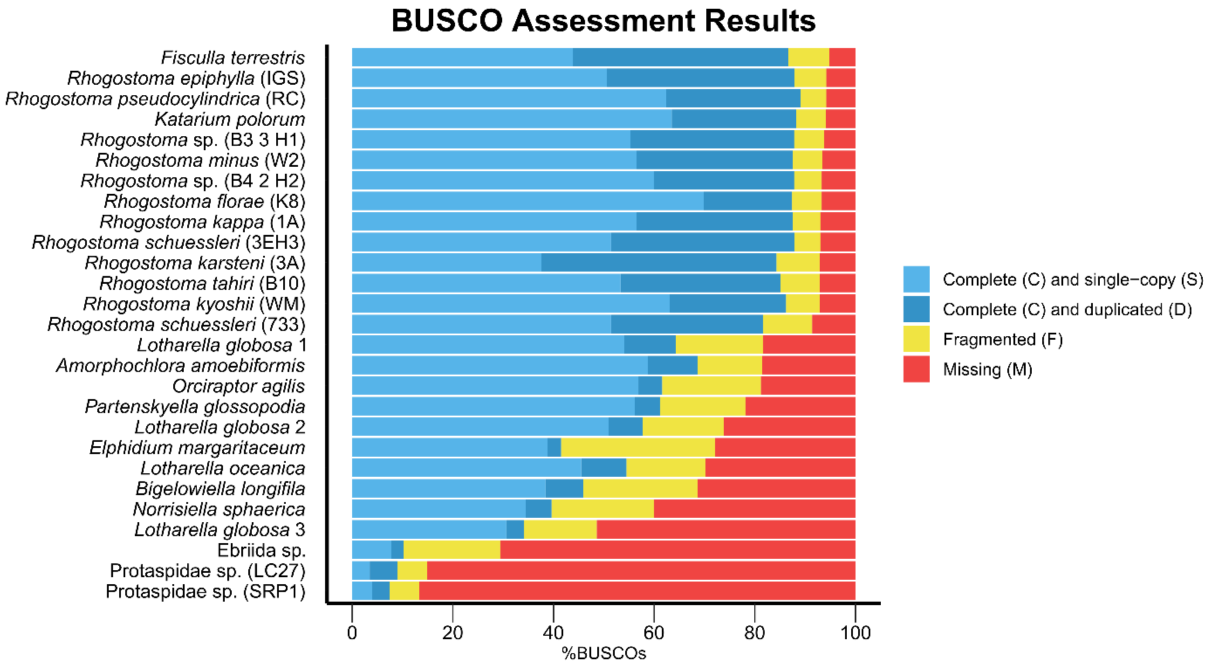


**Supplementary Figure 3: BUSCOs assessment of the Rhizaria transcriptomes.** BUSCO assessment of the Rhizaria transcriptomes based on the Eukaryota database. The colour code for complete single-copy orthologs (light blue), duplicated complete orthologs (dark blue), fragmented orthologs (yellow) and missing orthologs (red).

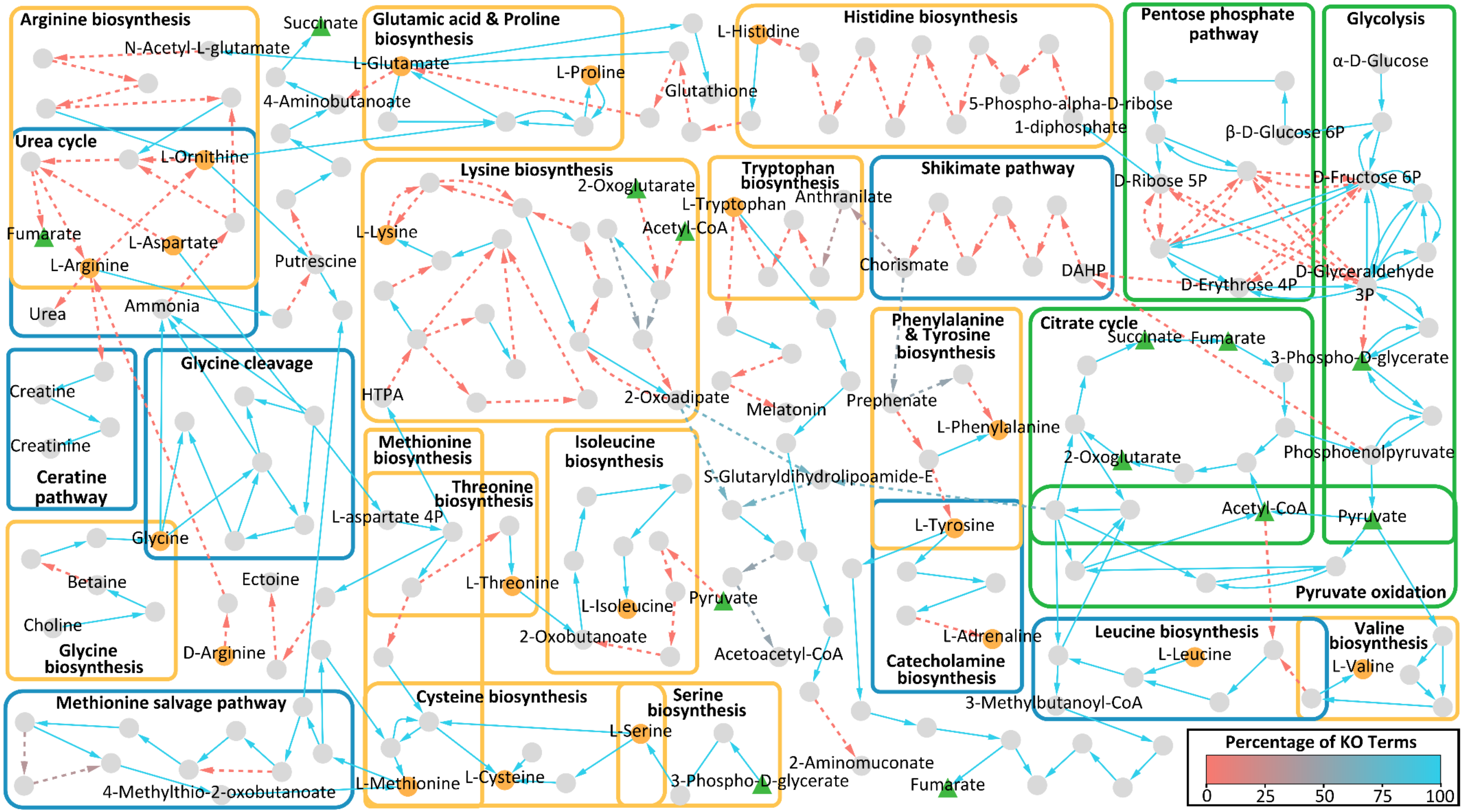


**Supplementary Figure 4: Overview of the central carbohydrate and amino acid metabolism for *Rhogostoma epiphylla* (IGS).** The graph illustrates a reconstruction of the central carbohydrate (green boxes) and amino acid metabolism for *Rhogostoma epiphylla* (IGS) based on KEGG ontologies and KEGG modules. Nodes represent components and edges represent enzymatic reactions. Amino acids are highlighted in orange, central compounds of carbohydrate metabolism in green. The edge color indicates the percentage of KO terms present, normalized to the minimum number of KO terms required for the respective reaction. Solid edges indicate that all KO terms were present for the respective reaction.

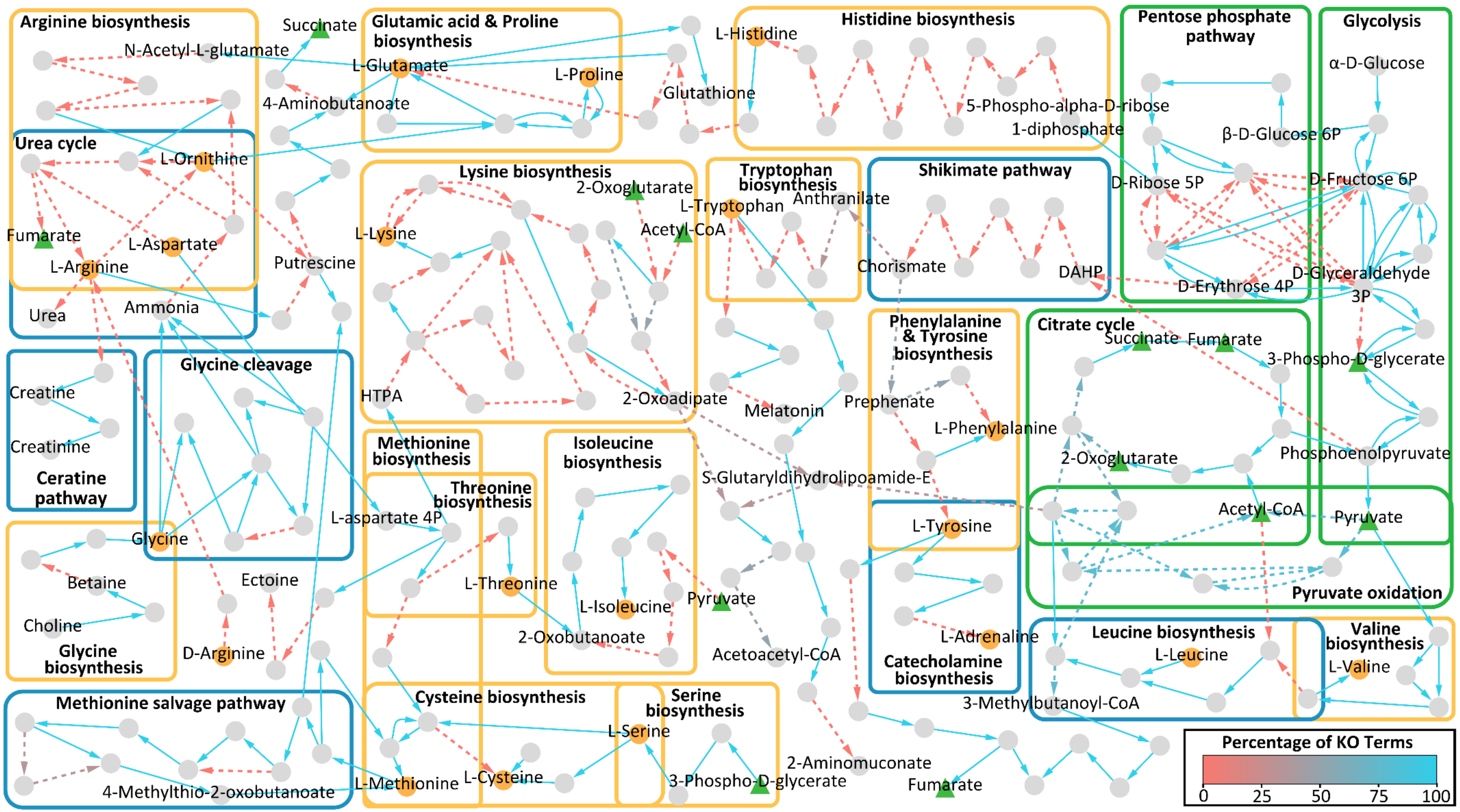


**Supplementary Figure 5: Overview of the central carbohydrate and amino acid metabolism for *Rhogostoma minus* (W2).** The graph illustrates a reconstruction of the central carbohydrate (green boxes) and amino acid metabolism for *Rhogostoma minus* (W2) based on KEGG ontologies and KEGG modules. Nodes represent components and edges represent enzymatic reactions. Amino acids are highlighted in orange, central compounds of carbohydrate metabolism in green. The edge color indicates the percentage of KO terms present, normalized to the minimum number of KO terms required for the respective reaction. Solid edges indicate that all KO terms were present for the respective reaction.

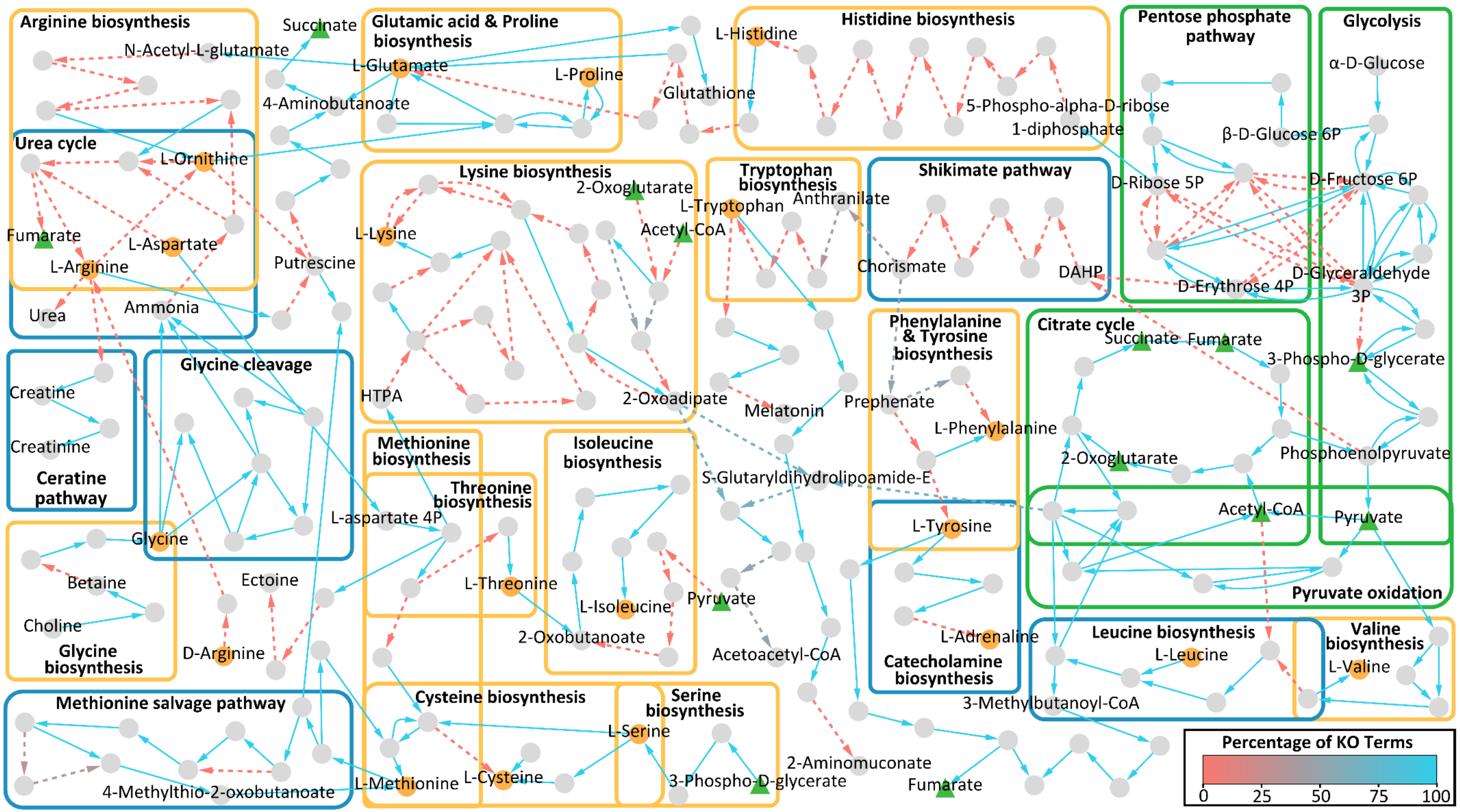


**Supplementary Figure 6: Overview of the central carbohydrate and amino acid metabolism for *Rhogostoma kappa* (1A).** The graph illustrates a reconstruction of the central carbohydrate (green boxes) and amino acid metabolism for *Rhogostoma kappa* (1A) based on KEGG ontologies and KEGG modules. Nodes represent components and edges represent enzymatic reactions. Amino acids are highlighted in orange, central compounds of carbohydrate metabolism in green. The edge color indicates the percentage of KO terms present, normalized to the minimum number of KO terms required for the respective reaction. Solid edges indicate that all KO terms were present for the respective reaction.

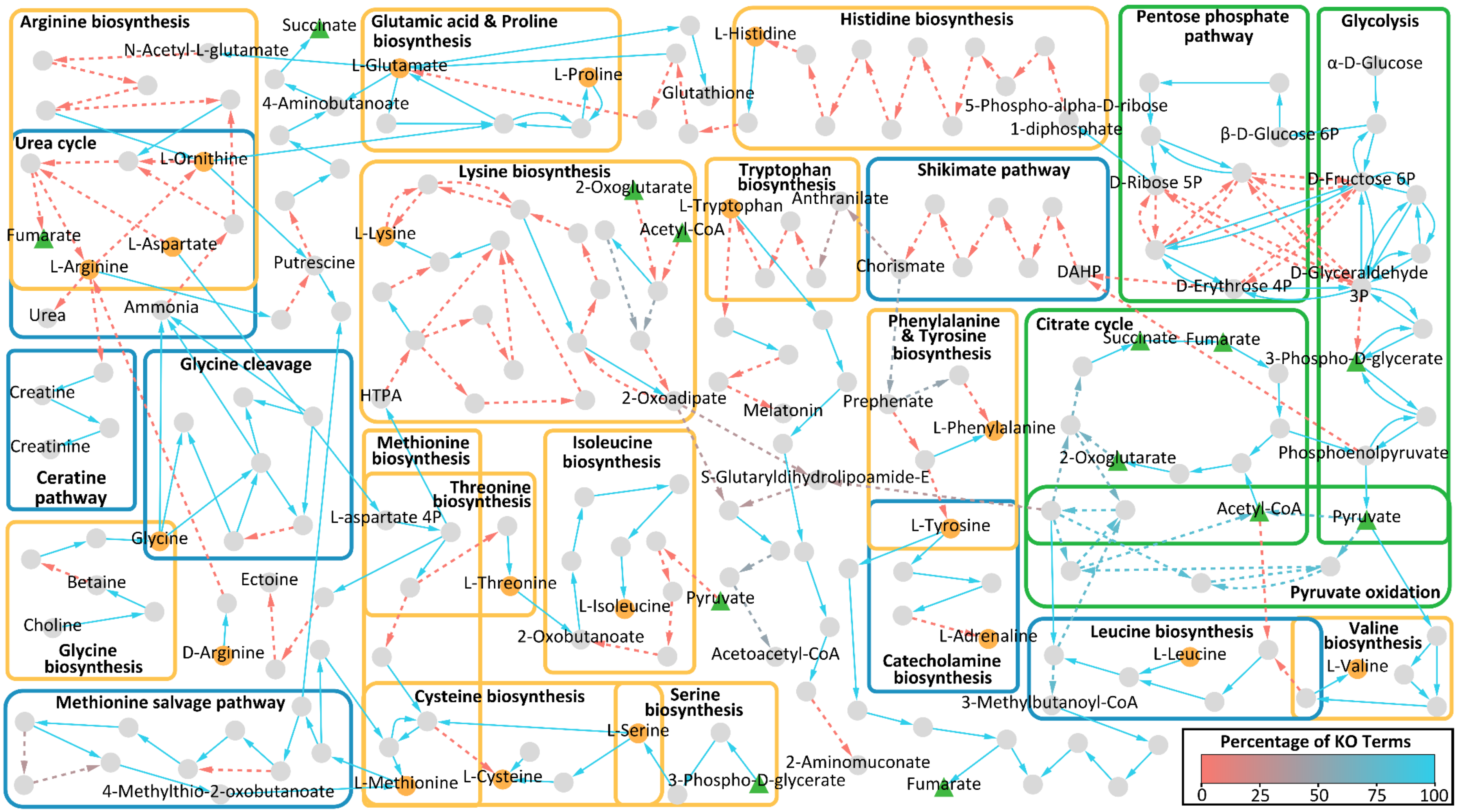


**Supplementary Figure 7: Overview of the central carbohydrate and amino acid metabolism for *Rhogostoma karsteni* (3A).** The graph illustrates a reconstruction of the central carbohydrate (green boxes) and amino acid metabolism for *Rhogostoma karsteni* (3A) based on KEGG ontologies and KEGG modules. Nodes represent components and edges represent enzymatic reactions. Amino acids are highlighted in orange, central compounds of carbohydrate metabolism in green. The edge color indicates the percentage of KO terms present, normalized to the minimum number of KO terms required for the respective reaction. Solid edges indicate that all KO terms were present for the respective reaction.

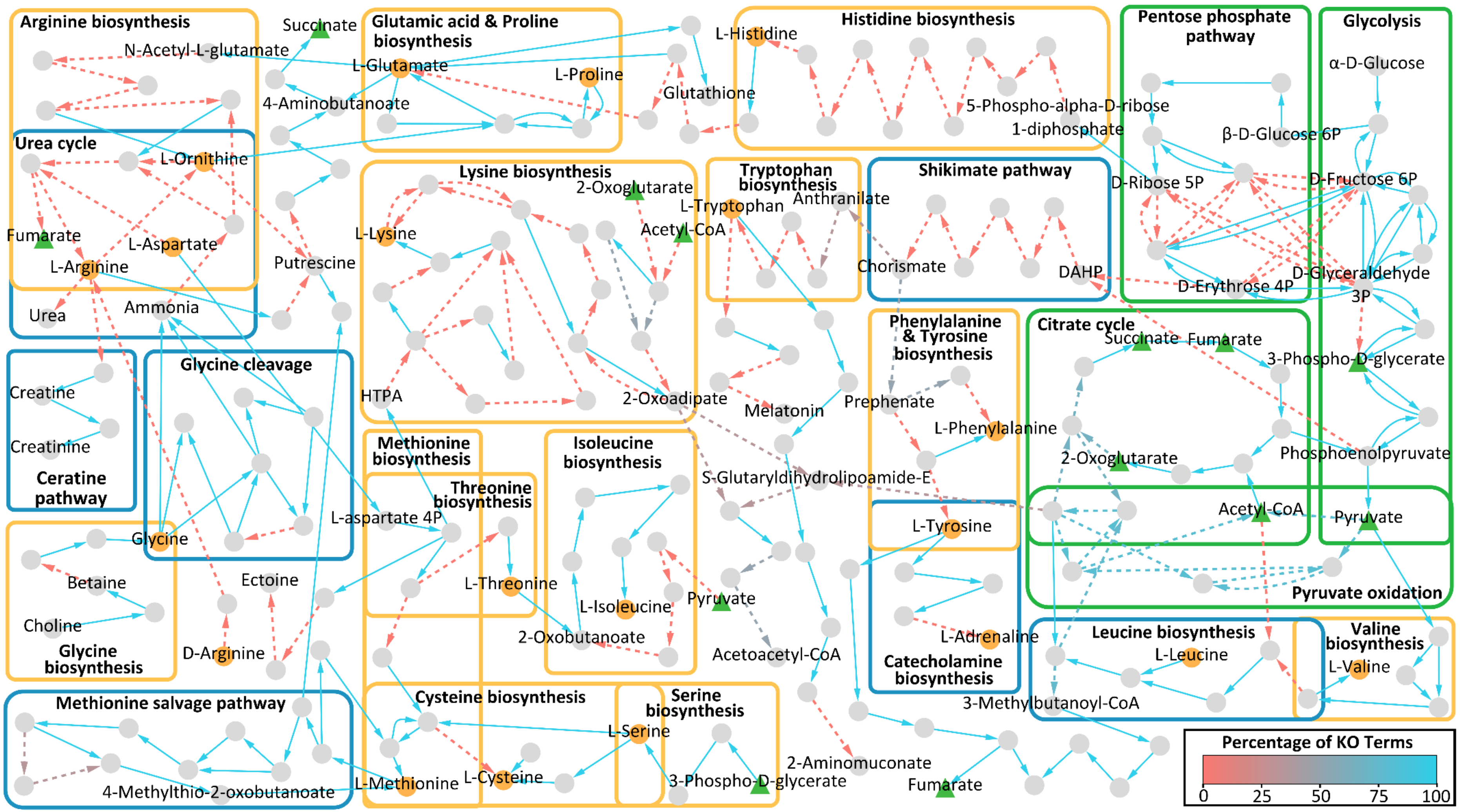


**Supplementary Figure 8: Overview of the central carbohydrate and amino acid metabolism for *Rhogostoma tahiri* (B10).** The graph illustrates a reconstruction of the central carbohydrate (green boxes) and amino acid metabolism for *Rhogostoma tahiri* (B10) based on KEGG ontologies and KEGG modules. Nodes represent components and edges represent enzymatic reactions. Amino acids are highlighted in orange, central compounds of carbohydrate metabolism in green. The edge color indicates the percentage of KO terms present, normalized to the minimum number of KO terms required for the respective reaction. Solid edges indicate that all KO terms were present for the respective reaction.

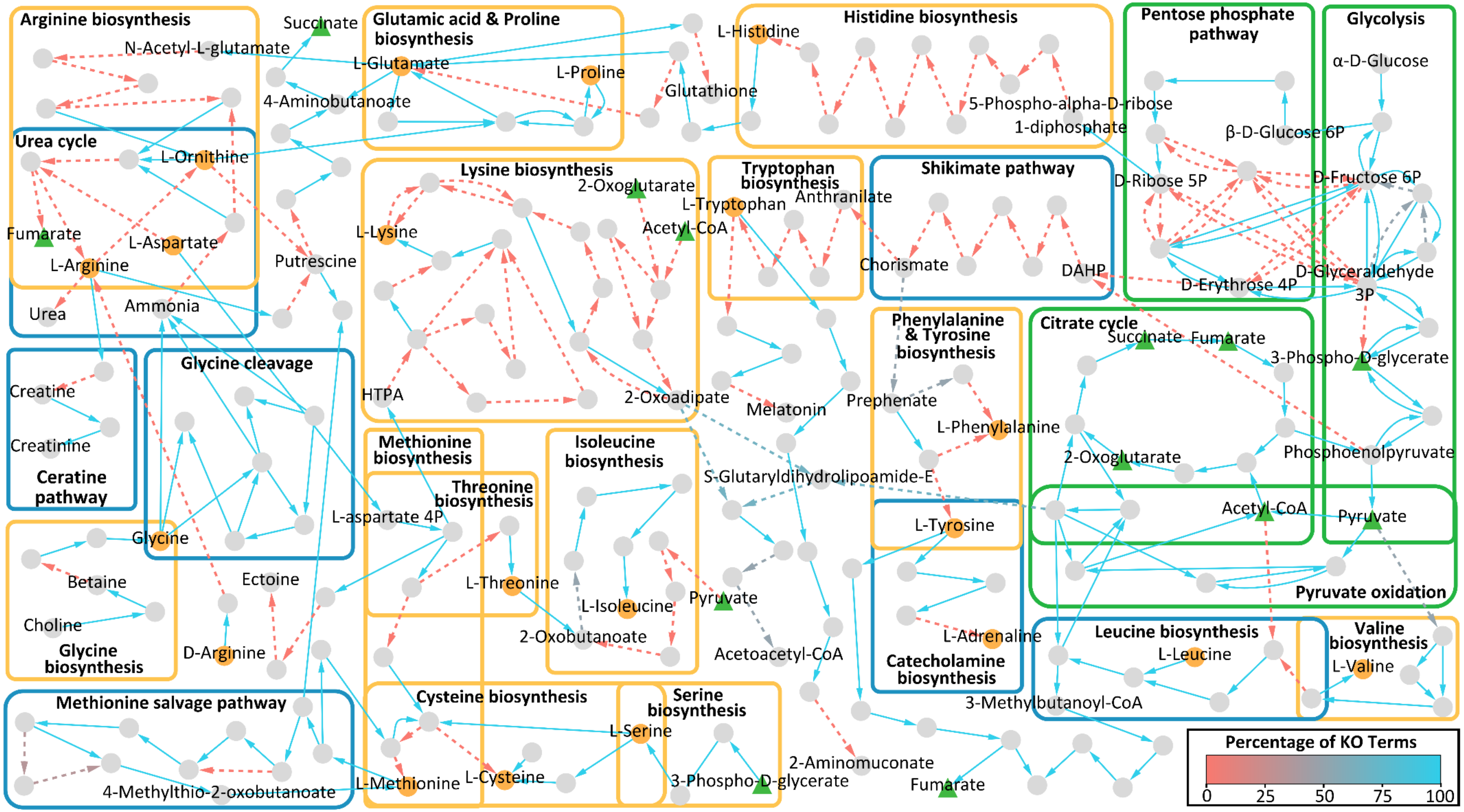


**Supplementary Figure 9: Overview of the central carbohydrate and amino acid metabolism for *Rhogostoma florae* (K8).** The graph illustrates a reconstruction of the central carbohydrate (green boxes) and amino acid metabolism for *Rhogostoma florae* (K8) based on KEGG ontologies and KEGG modules. Nodes represent components and edges represent enzymatic reactions. Amino acids are highlighted in orange, central compounds of carbohydrate metabolism in green. The edge color indicates the percentage of KO terms present, normalized to the minimum number of KO terms required for the respective reaction. Solid edges indicate that all KO terms were present for the respective reaction.

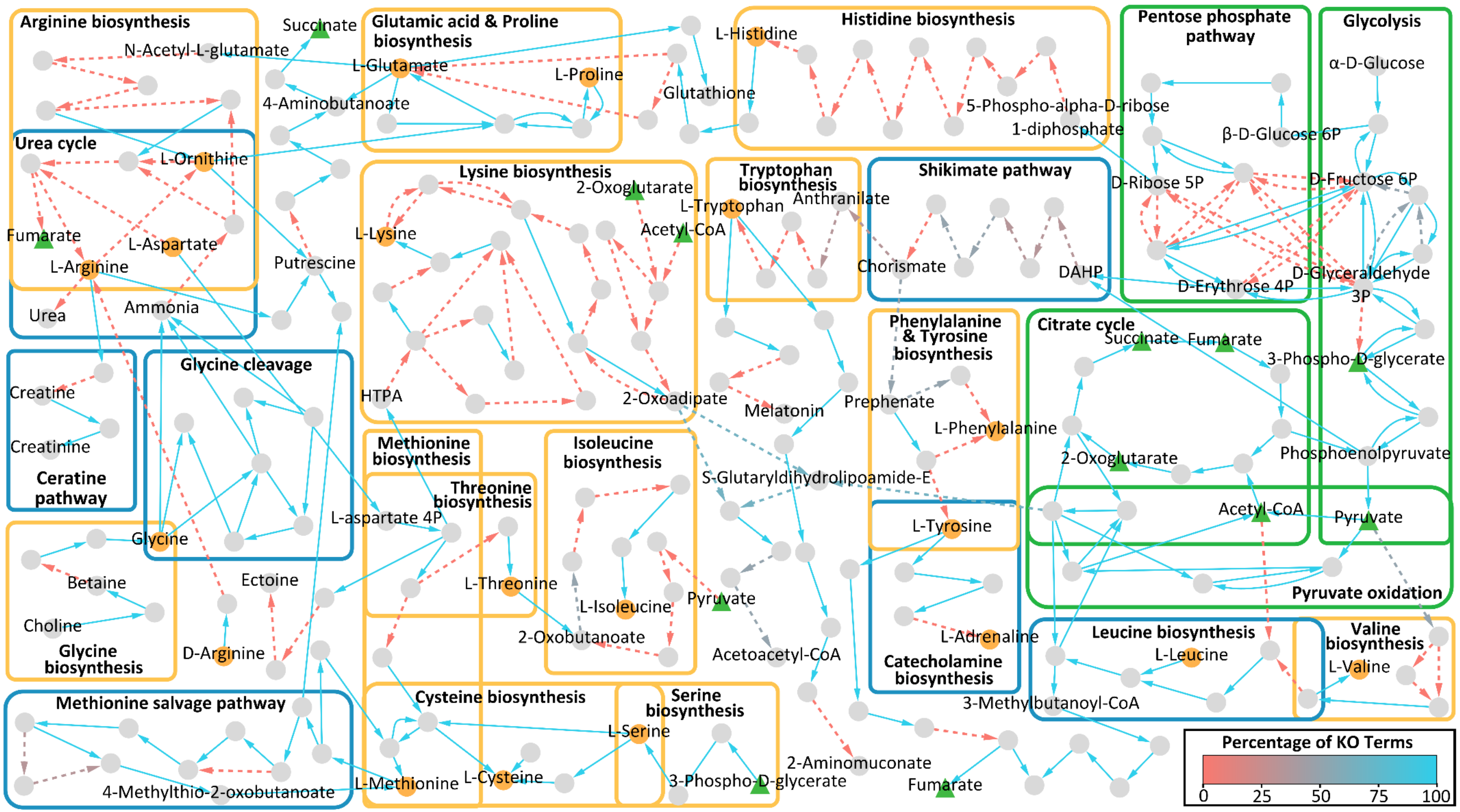


**Supplementary Figure 10: Overview of the central carbohydrate and amino acid metabolism for *Rhogostoma pseudocylindrica* (RC).** The graph illustrates a reconstruction of the central carbohydrate (green boxes) and amino acid metabolism for *Rhogostoma pseudocylindrica* (RC) based on KEGG ontologies and KEGG modules. Nodes represent components and edges represent enzymatic reactions. Amino acids are highlighted in orange, central compounds of carbohydrate metabolism in green. The edge color indicates the percentage of KO terms present, normalized to the minimum number of KO terms required for the respective reaction. Solid edges indicate that all KO terms were present for the respective reaction.

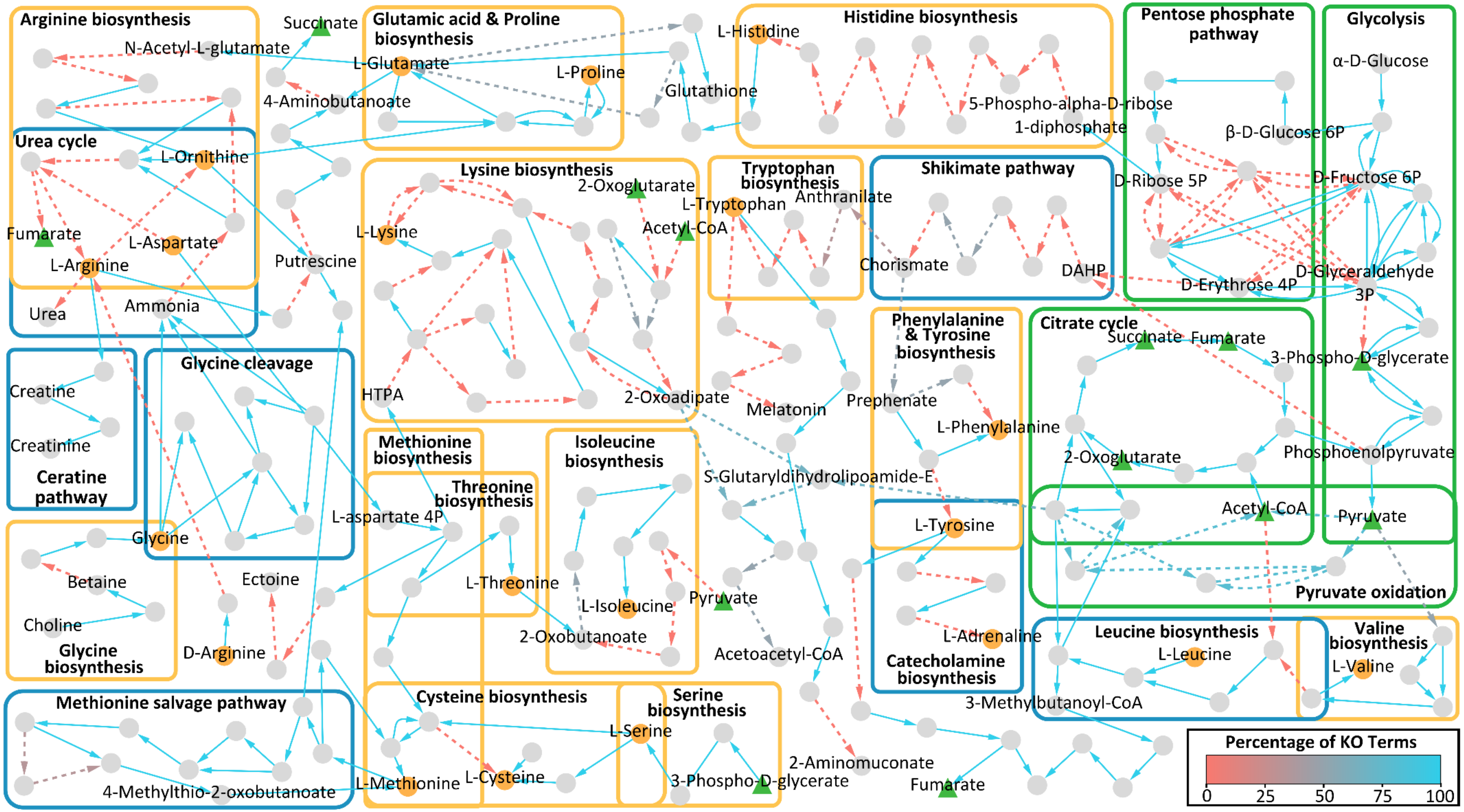


**Supplementary Figure 11: Overview of the central carbohydrate and amino acid metabolism for *Rhogostoma* sp. (B3 3 H1).** The graph illustrates a reconstruction of the central carbohydrate (green boxes) and amino acid metabolism for *Rhogostoma* sp. (B3 3 H1) based on KEGG ontologies and KEGG modules. Nodes represent components and edges represent enzymatic reactions. Amino acids are highlighted in orange, central compounds of carbohydrate metabolism in green. The edge color indicates the percentage of KO terms present, normalized to the minimum number of KO terms required for the respective reaction. Solid edges indicate that all KO terms were present for the respective reaction.

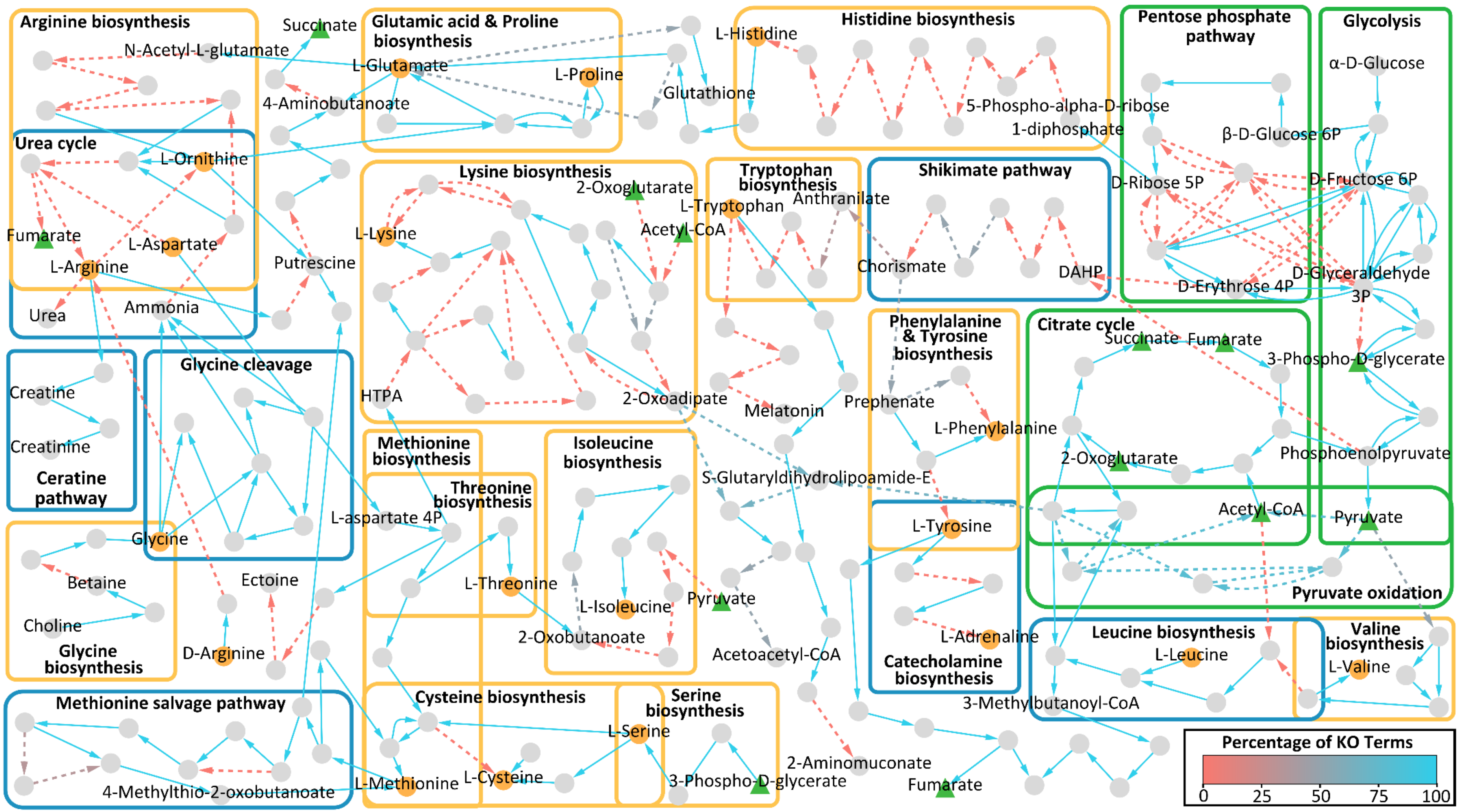


**Supplementary Figure 12: Overview of the central carbohydrate and amino acid metabolism for *Rhogostoma* sp. (B4 2 H2).** The graph illustrates a reconstruction of the central carbohydrate (green boxes) and amino acid metabolism for *Rhogostoma* sp. (B4 2 H2) based on KEGG ontologies and KEGG modules. Nodes represent components and edges represent enzymatic reactions. Amino acids are highlighted in orange, central compounds of carbohydrate metabolism in green. The edge color indicates the percentage of KO terms present, normalized to the minimum number of KO terms required for the respective reaction. Solid edges indicate that all KO terms were present for the respective reaction.

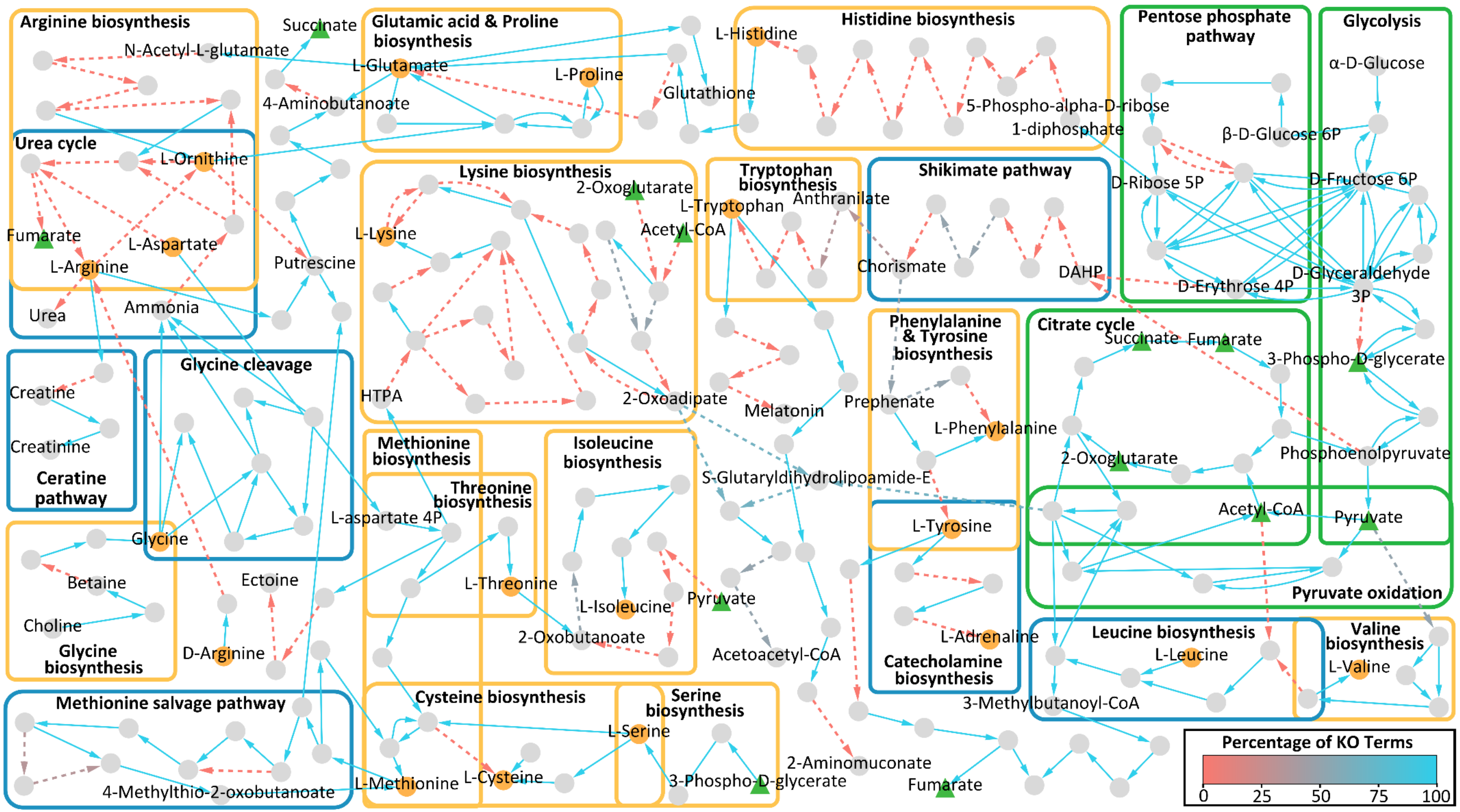


**Supplementary Figure 13: Overview of the central carbohydrate and amino acid metabolism for *Rhogostoma schuessleri* (733).** The graph illustrates a reconstruction of the central carbohydrate (green boxes) and amino acid metabolism for *Rhogostoma schuessleri* (733) based on KEGG ontologies and KEGG modules. Nodes represent components and edges represent enzymatic reactions. Amino acids are highlighted in orange, central compounds of carbohydrate metabolism in green. The edge color indicates the percentage of KO terms present, normalized to the minimum number of KO terms required for the respective reaction. Solid edges indicate that all KO terms were present for the respective reaction.

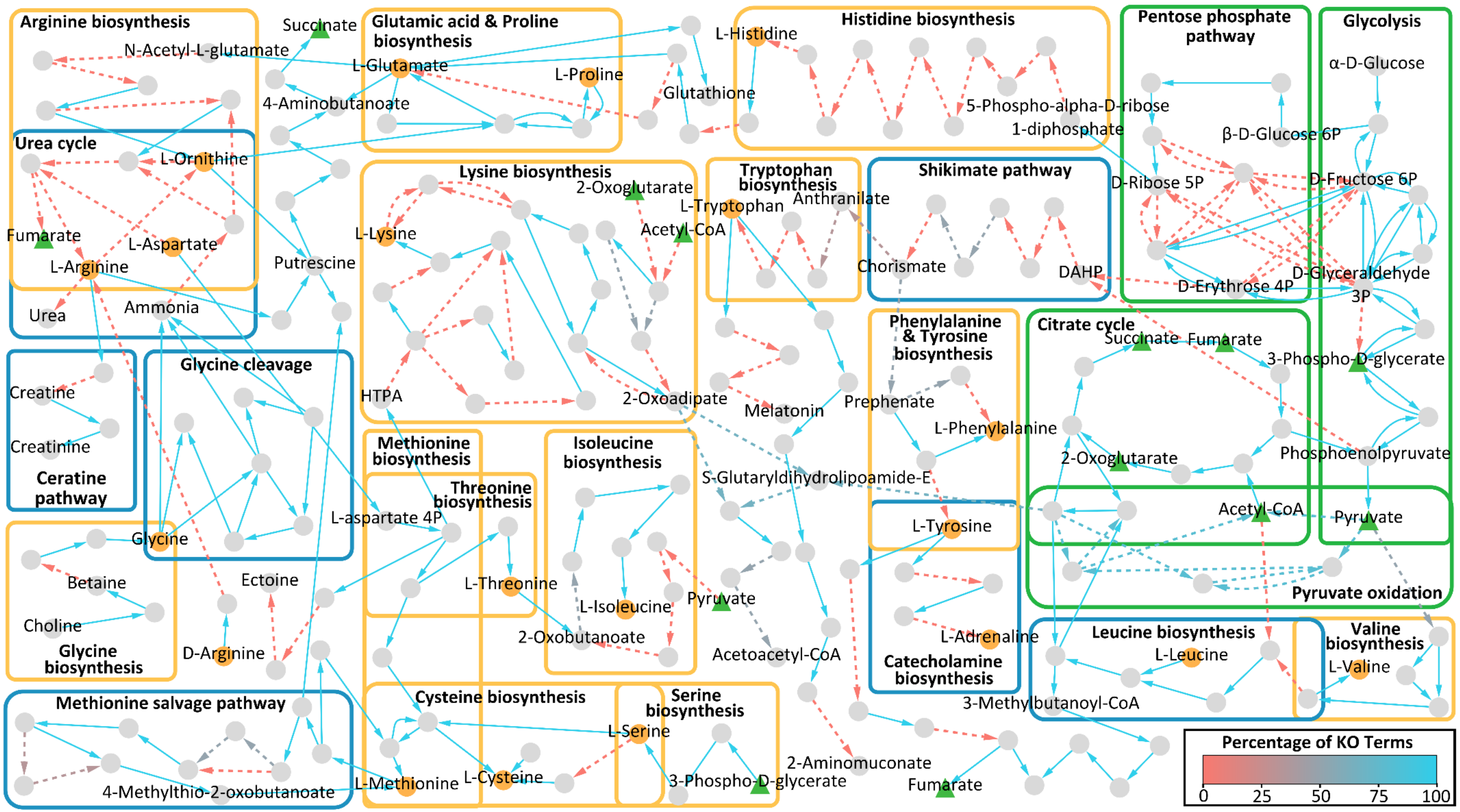


**Supplementary Figure 14: Overview of the central carbohydrate and amino acid metabolism for *Rhogostoma schuessleri* (3EH3).** The graph illustrates a reconstruction of the central carbohydrate (green boxes) and amino acid metabolism for *Rhogostoma schuessleri* (3EH3) based on KEGG ontologies and KEGG modules. Nodes represent components and edges represent enzymatic reactions. Amino acids are highlighted in orange, central compounds of carbohydrate metabolism in green. The edge color indicates the percentage of KO terms present, normalized to the minimum number of KO terms required for the respective reaction. Solid edges indicate that all KO terms were present for the respective reaction.

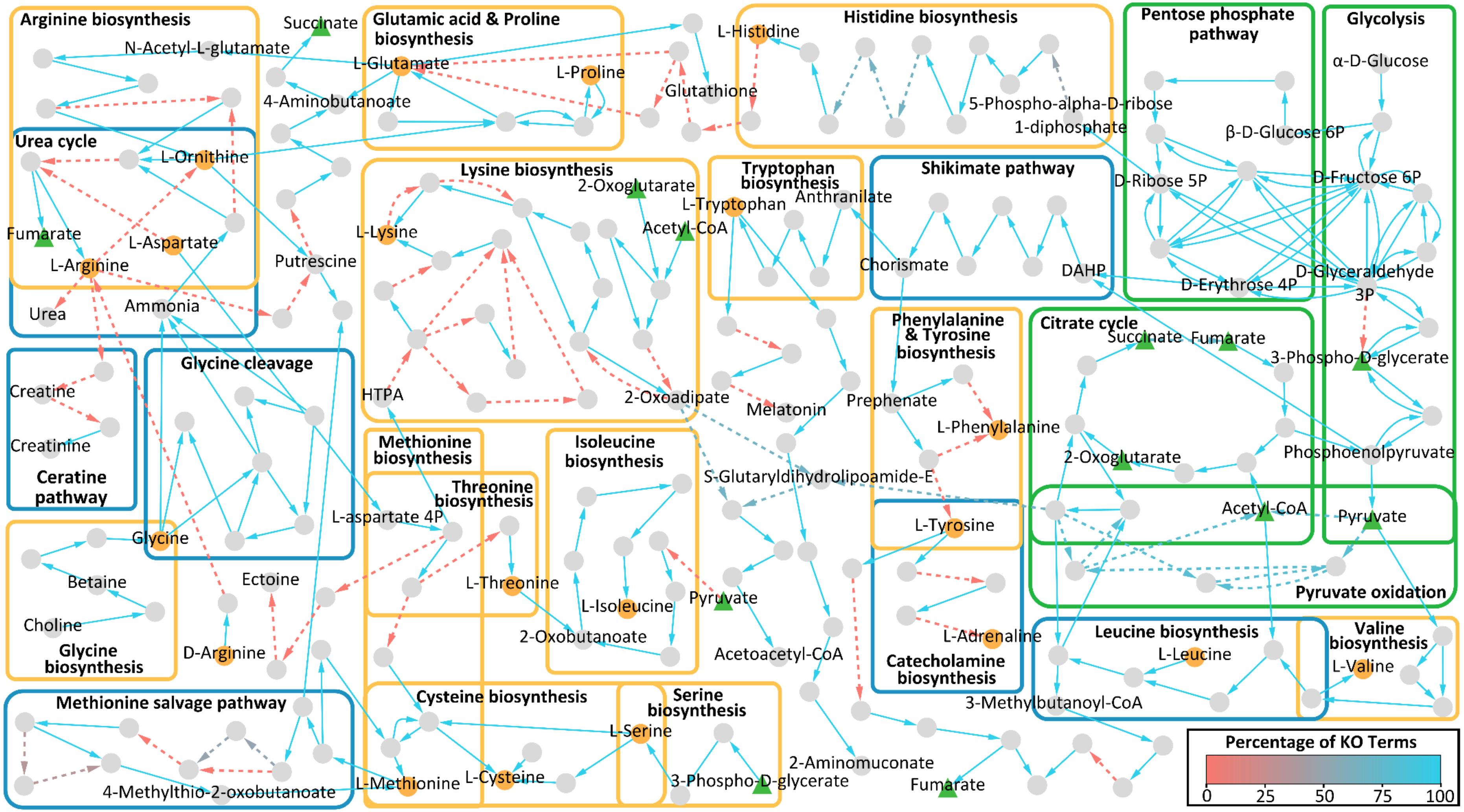


**Supplementary Figure 15: Overview of the central carbohydrate and amino acid metabolism for *Fisculla terrestris*.** The graph illustrates a reconstruction of the central carbohydrate (green boxes) and amino acid metabolism for *Fisculla terrestris* based on KEGG ontologies and KEGG modules. Nodes represent components and edges represent enzymatic reactions. Amino acids are highlighted in orange, central compounds of carbohydrate metabolism in green. The edge color indicates the percentage of KO terms present, normalized to the minimum number of KO terms required for the respective reaction. Solid edges indicate that all KO terms were present for the respective reaction.

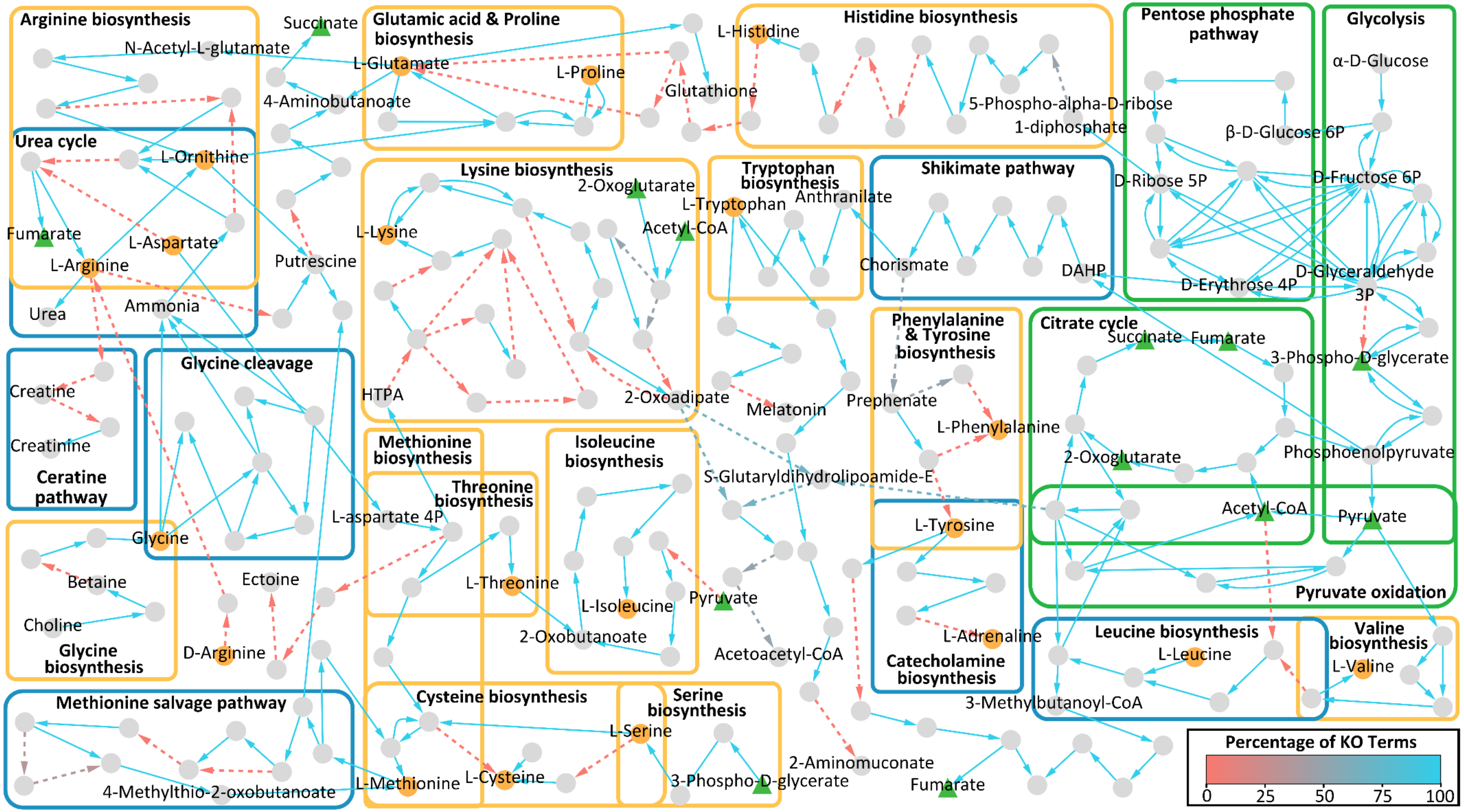


**Supplementary Figure 16: Overview of the central carbohydrate and amino acid metabolism for *Katarium polorum*.** The graph illustrates a reconstruction of the central carbohydrate (green boxes) and amino acid metabolism for *Katarium polorum* based on KEGG ontologies and KEGG modules. Nodes represent components and edges represent enzymatic reactions. Amino acids are highlighted in orange, central compounds of carbohydrate metabolism in green. The edge color indicates the percentage of KO terms present, normalized to the minimum number of KO terms required for the respective reaction. Solid edges indicate that all KO terms were present for the respective reaction.

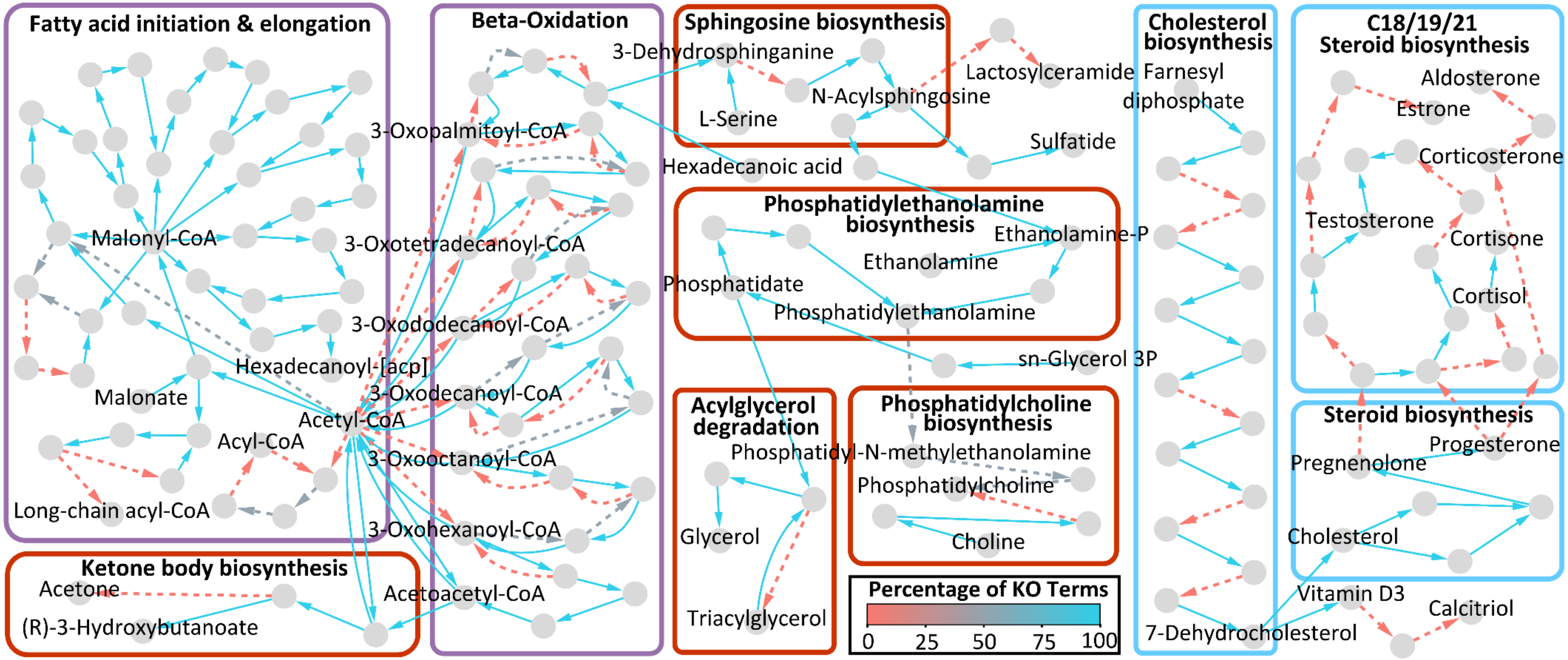


**Supplementary Figure 17: Overview of the lipid metabolism for *Rhogostoma epiphylla* (IGS) .** The graph illustrates a reconstruction of the fatty acid (purple boxes), sterol (blue boxes), and lipid (red boxes) metabolism for *Rhogostoma epiphylla* (IGS) based on KEGG ontologies and KEGG modules. Nodes represent components and edges represent enzymatic reactions. The edge color indicates the percentage of KO terms present, normalized to the minimum number of KO terms required for the respective reaction. Solid edges indicate that all KO terms were present for the respective reaction.

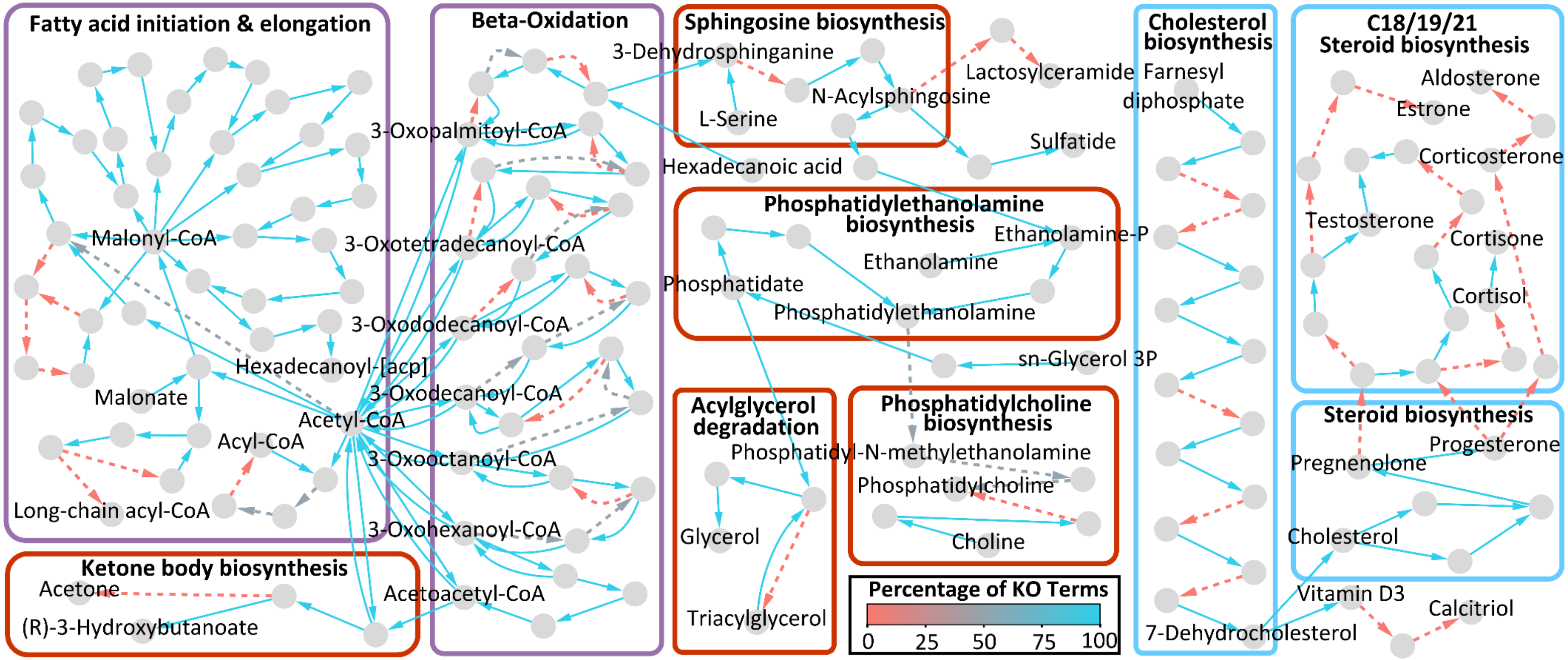


**Supplementary Figure 18: Overview of the lipid metabolism for *Rhogostoma minus* (W2).** The graph illustrates a reconstruction of the fatty acid (purple boxes), sterol (blue boxes), and lipid (red boxes) metabolism for *Rhogostoma minus* (W2) based on KEGG ontologies and KEGG modules. Nodes represent components and edges represent enzymatic reactions. The edge color indicates the percentage of KO terms present, normalized to the minimum number of KO terms required for the respective reaction. Solid edges indicate that all KO terms were present for the respective reaction.

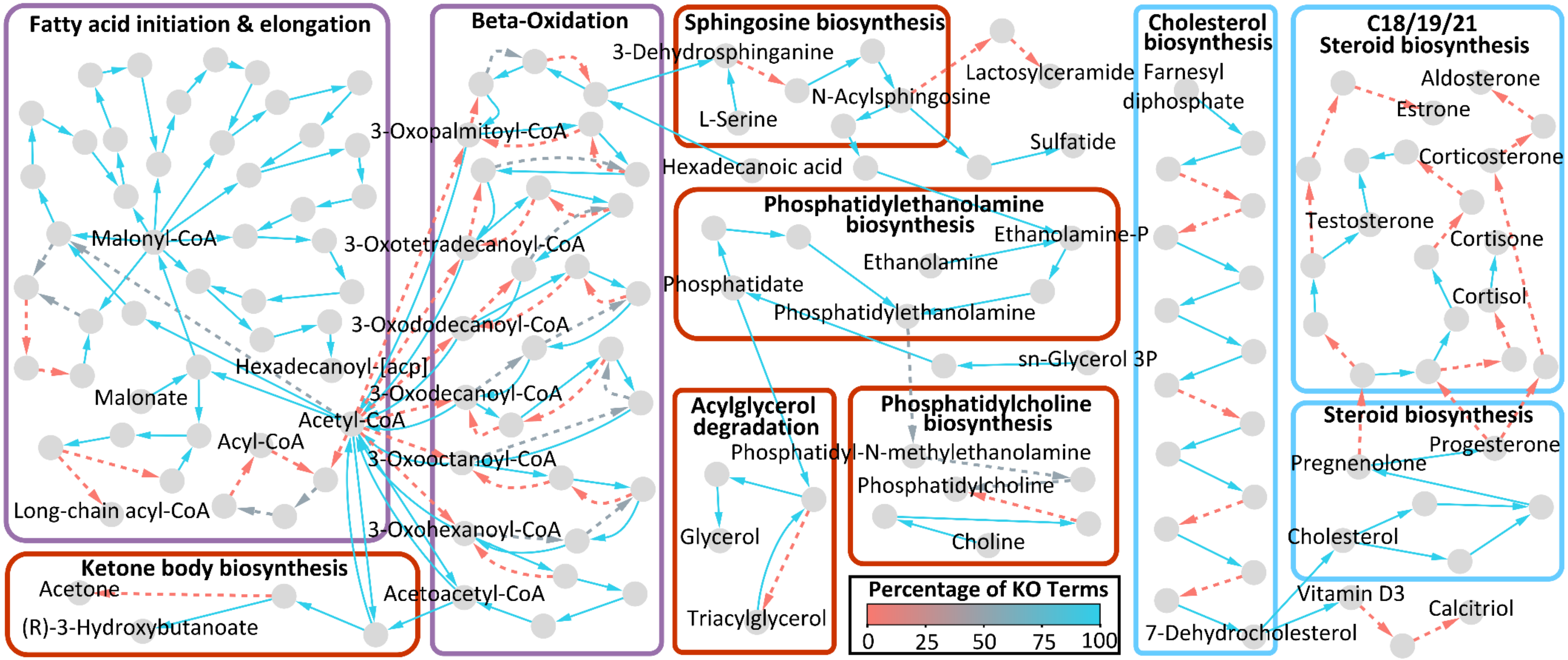


**Supplementary Figure 19: Overview of the lipid metabolism for *Rhogostoma kappa* (1A).** The graph illustrates a reconstruction of the fatty acid (purple boxes), sterol (blue boxes), and lipid (red boxes) metabolism for *Rhogostoma kappa* (1A) based on KEGG ontologies and KEGG modules. Nodes represent components and edges represent enzymatic reactions. The edge color indicates the percentage of KO terms present, normalized to the minimum number of KO terms required for the respective reaction. Solid edges indicate that all KO terms were present for the respective reaction.

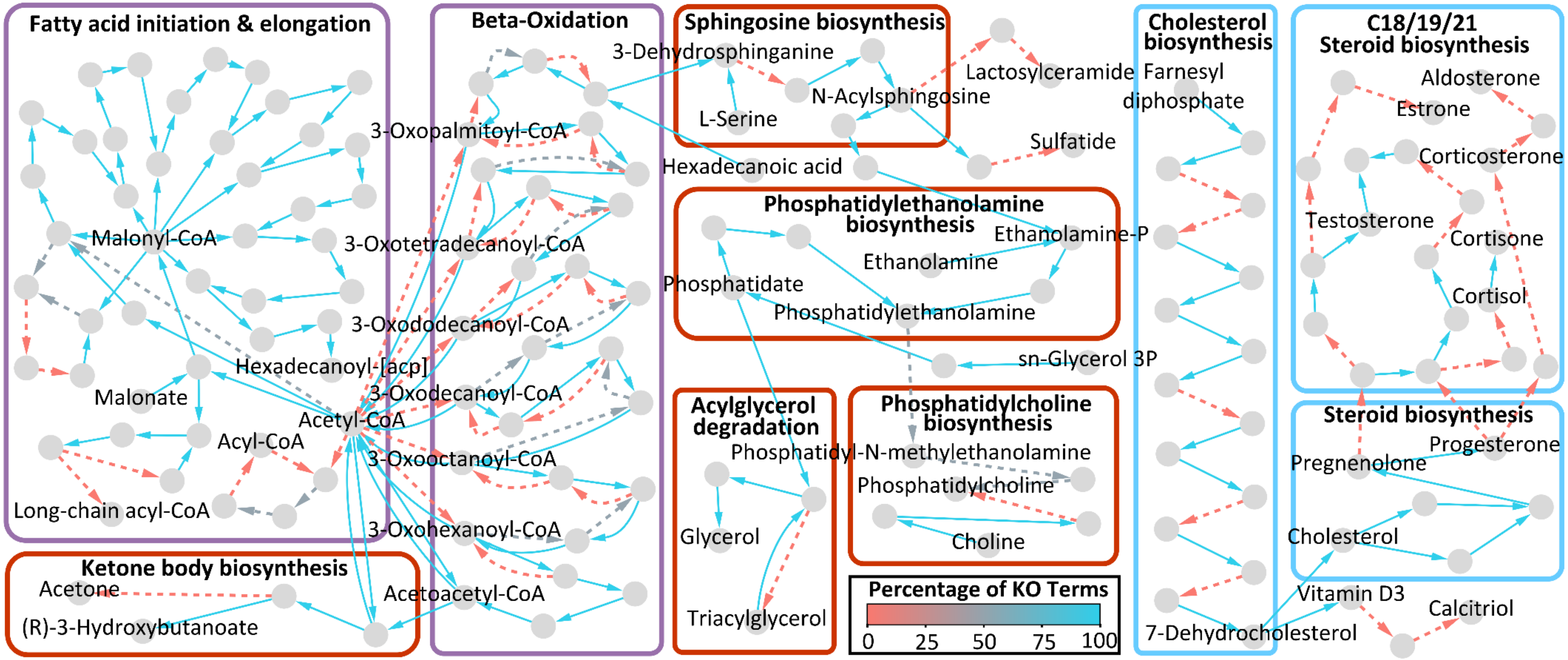


**Supplementary Figure 20: Overview of the lipid metabolism for *Rhogostoma karsteni* (3A).** The graph illustrates a reconstruction of the fatty acid (purple boxes), sterol (blue boxes), and lipid (red boxes) metabolism for *Rhogostoma karsteni* (3A) based on KEGG ontologies and KEGG modules. Nodes represent components and edges represent enzymatic reactions. The edge color indicates the percentage of KO terms present, normalized to the minimum number of KO terms required for the respective reaction. Solid edges indicate that all KO terms were present for the respective reaction.

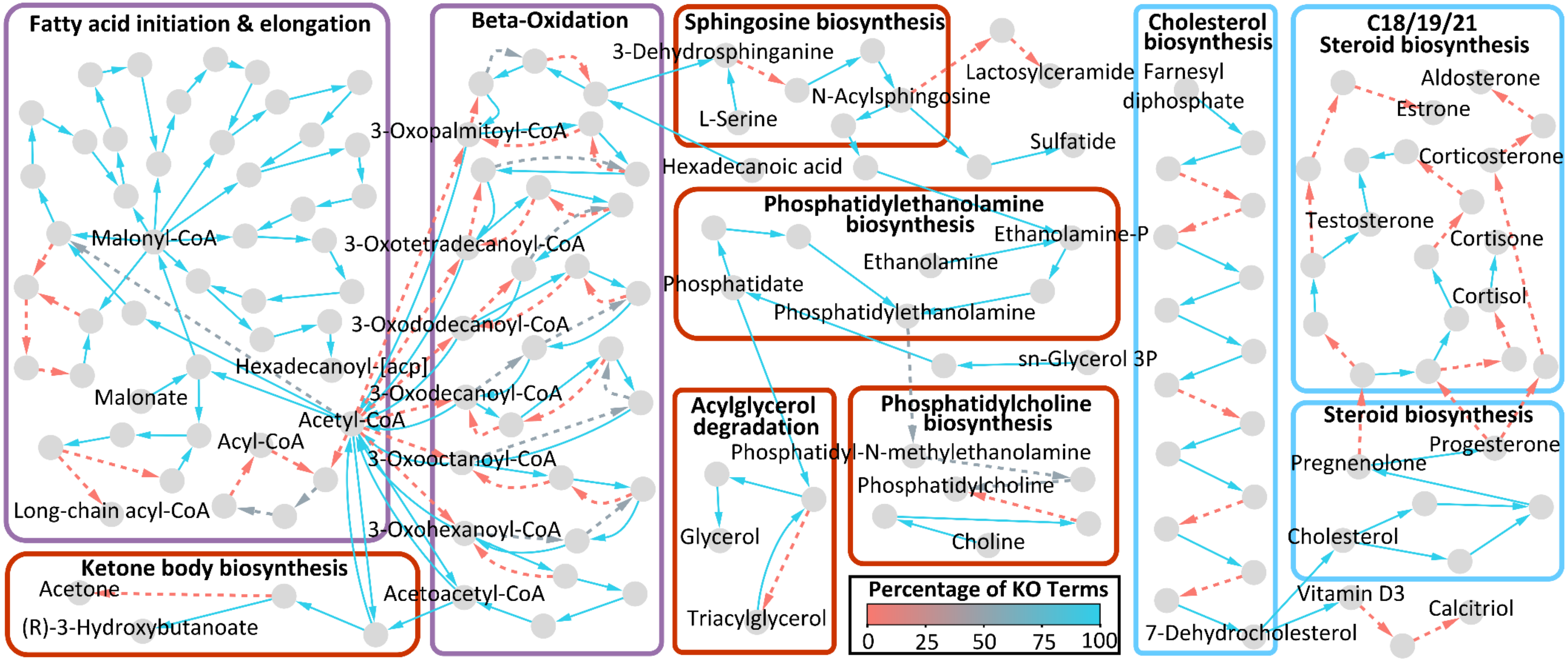


**Supplementary Figure 21: Overview of the lipid metabolism for *Rhogostoma tahiri* (B10).** The graph illustrates a reconstruction of the fatty acid (purple boxes), sterol (blue boxes), and lipid (red boxes) metabolism for *Rhogostoma tahiri* (B10) based on KEGG ontologies and KEGG modules. Nodes represent components and edges represent enzymatic reactions. The edge color indicates the percentage of KO terms present, normalized to the minimum number of KO terms required for the respective reaction. Solid edges indicate that all KO terms were present for the respective reaction.

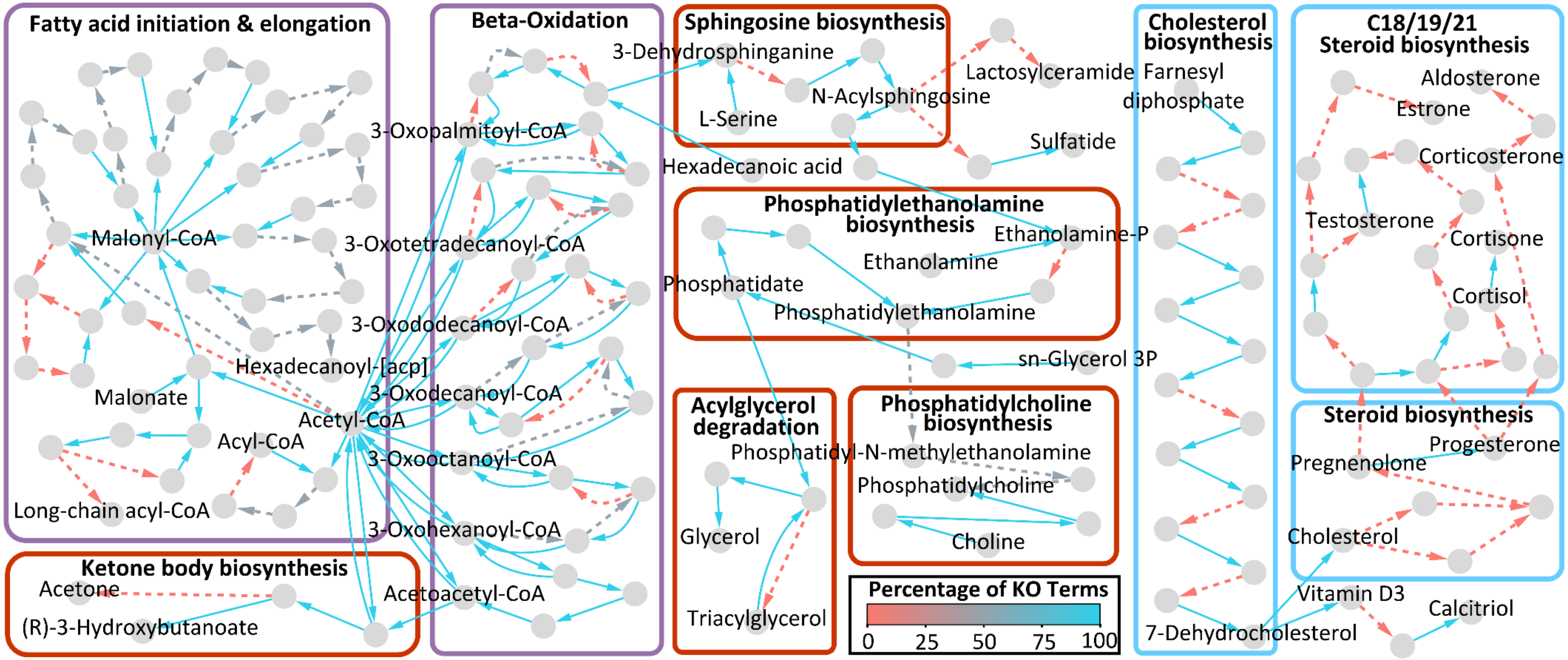


**Supplementary Figure 22: Overview of the lipid metabolism for *Rhogostoma florae* (K8).** The graph illustrates a reconstruction of the fatty acid (purple boxes), sterol (blue boxes), and lipid (red boxes) metabolism for *Rhogostoma florae* (K8) based on KEGG ontologies and KEGG modules. Nodes represent components and edges represent enzymatic reactions. The edge color indicates the percentage of KO terms present, normalized to the minimum number of KO terms required for the respective reaction. Solid edges indicate that all KO terms were present for the respective reaction.

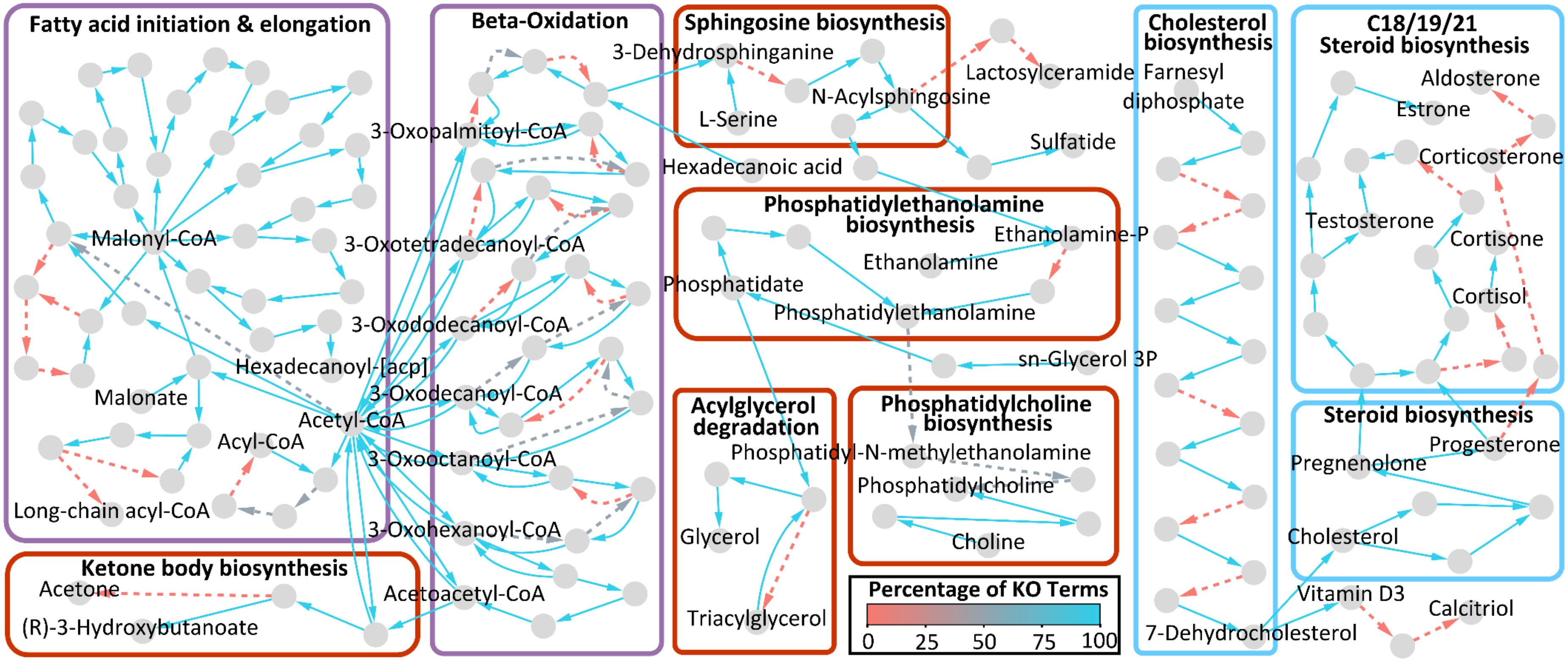


**Supplementary Figure 23: Overview of the lipid metabolism for *Rhogostoma pseudocylindrica* (RC).** The graph illustrates a reconstruction of the fatty acid (purple boxes), sterol (blue boxes), and lipid (red boxes) metabolism for *Rhogostoma pseudocylindrica* (RC) based on KEGG ontologies and KEGG modules. Nodes represent components and edges represent enzymatic reactions. The edge color indicates the percentage of KO terms present, normalized to the minimum number of KO terms required for the respective reaction. Solid edges indicate that all KO terms were present for the respective reaction.

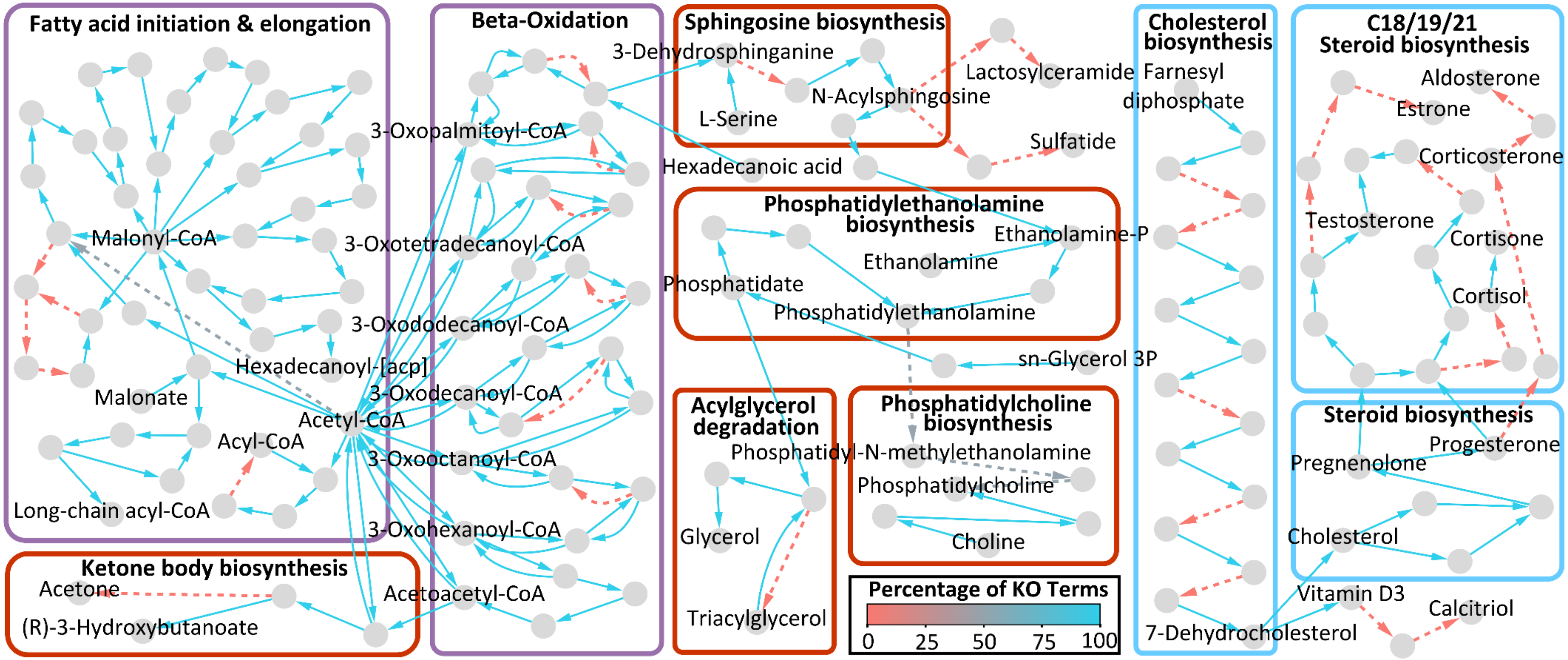


**Supplementary Figure 24: Overview of the lipid metabolism for *Rhogostoma* sp. (B3 3 H1).** The graph illustrates a reconstruction of the fatty acid (purple boxes), sterol (blue boxes), and lipid (red boxes) metabolism for *Rhogostoma* sp. (B3 3 H1) based on KEGG ontologies and KEGG modules. Nodes represent components and edges represent enzymatic reactions. The edge color indicates the percentage of KO terms present, normalized to the minimum number of KO terms required for the respective reaction. Solid edges indicate that all KO terms were present for the respective reaction.

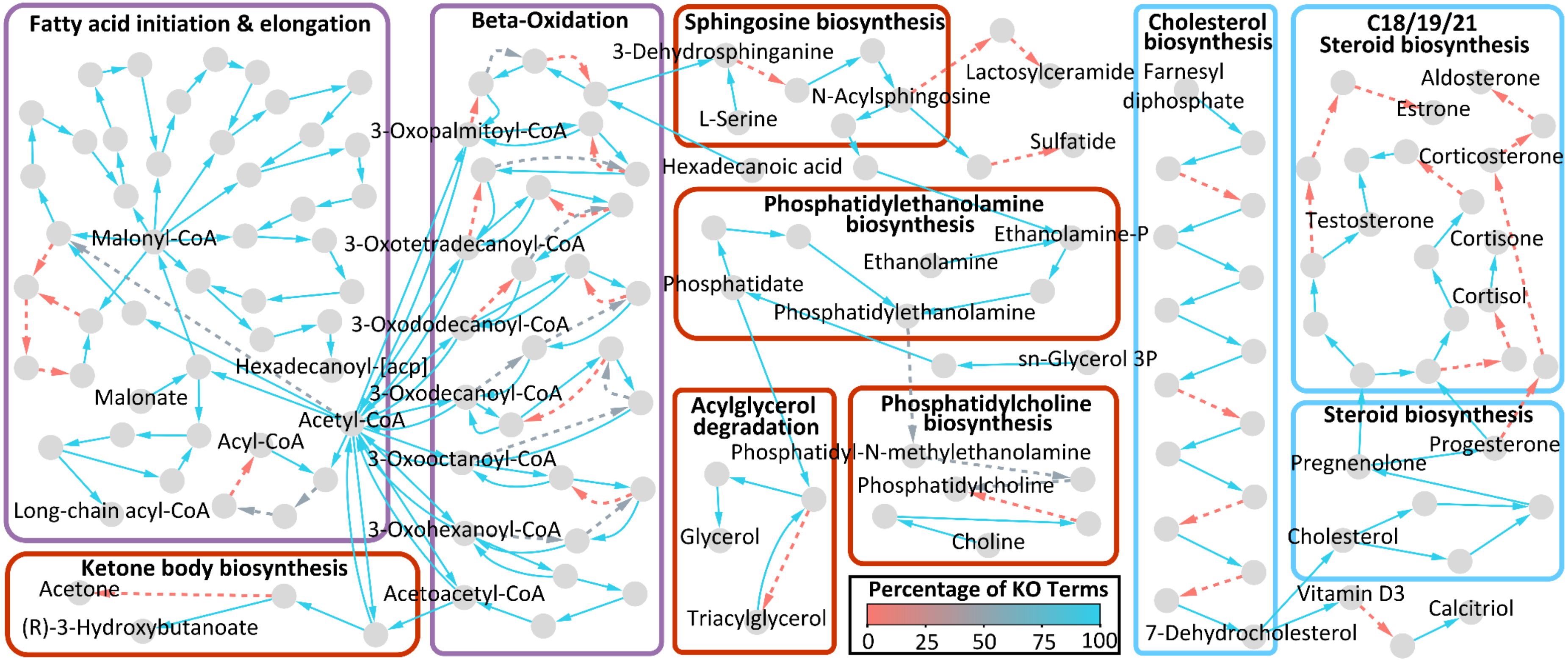


**Supplementary Figure 25: Overview of the lipid metabolism for *Rhogostoma* sp. (B4 2 H2).** The graph illustrates a reconstruction of the fatty acid (purple boxes), sterol (blue boxes), and lipid (red boxes) metabolism for *Rhogostoma* sp. (B4 2 H2) based on KEGG ontologies and KEGG modules. Nodes represent components and edges represent enzymatic reactions. The edge color indicates the percentage of KO terms present, normalized to the minimum number of KO terms required for the respective reaction. Solid edges indicate that all KO terms were present for the respective reaction.

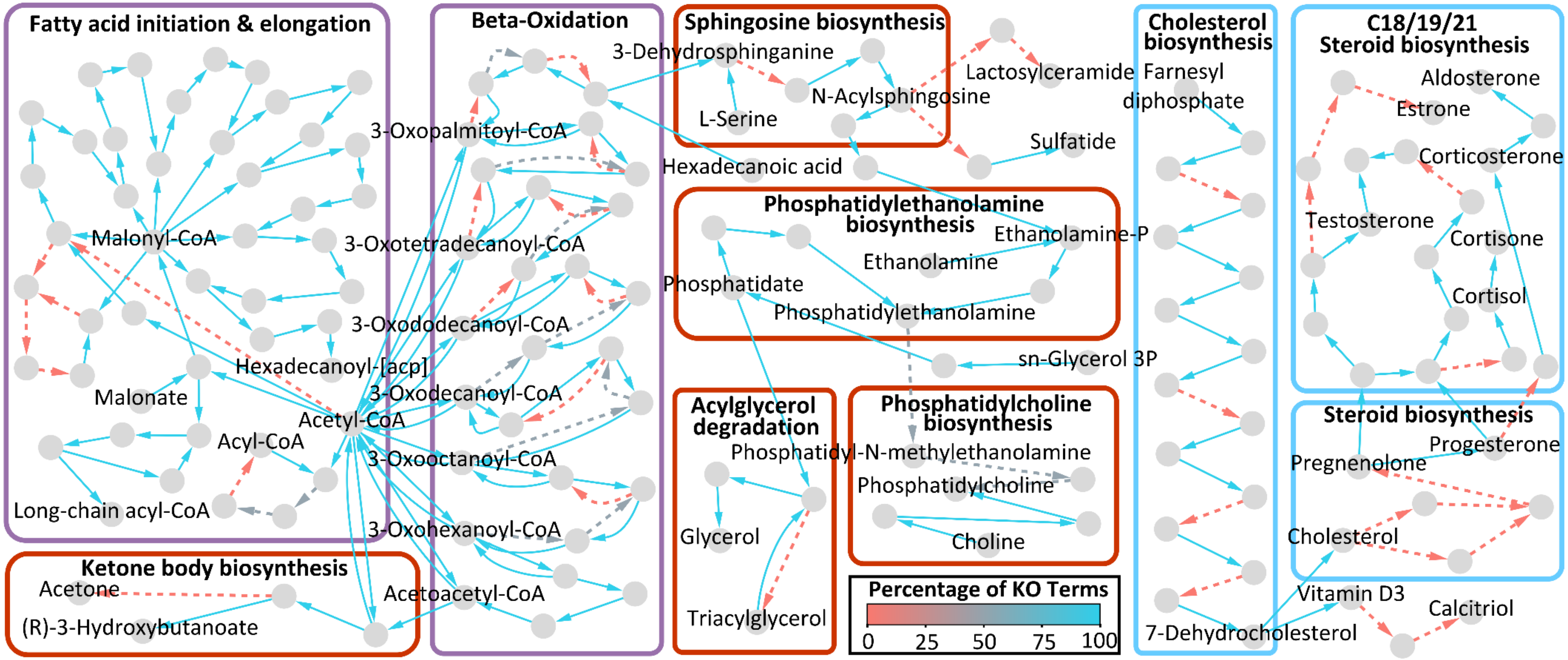


**Supplementary Figure 26: Overview of the lipid metabolism for *Rhogostoma schuessleri* (733).** The graph illustrates a reconstruction of the fatty acid (purple boxes), sterol (blue boxes), and lipid (red boxes) metabolism for *Rhogostoma schuessleri* (733) based on KEGG ontologies and KEGG modules. Nodes represent components and edges represent enzymatic reactions. The edge color indicates the percentage of KO terms present, normalized to the minimum number of KO terms required for the respective reaction. Solid edges indicate that all KO terms were present for the respective reaction.

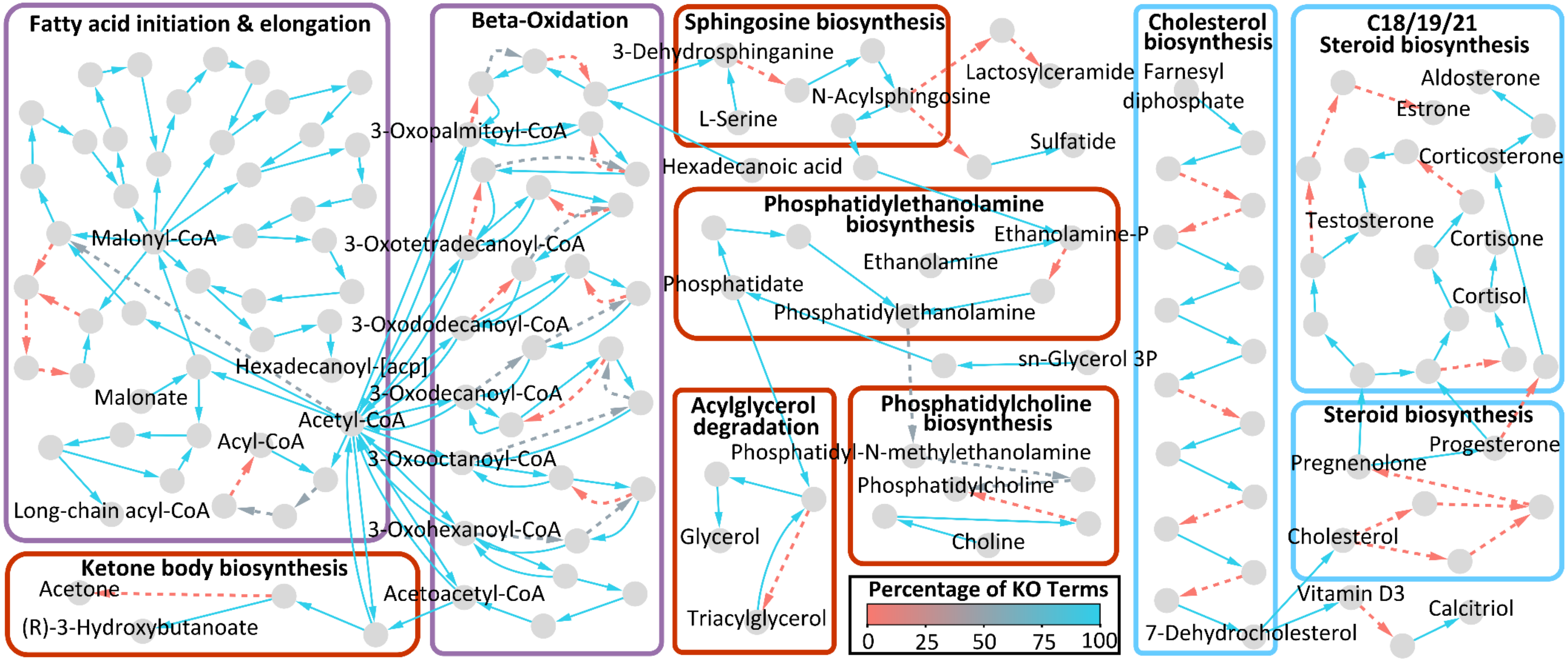


**Supplementary Figure 27: Overview of the lipid metabolism for *Rhogostoma schuessleri* (3EH3).** The graph illustrates a reconstruction of the fatty acid (purple boxes), sterol (blue boxes), and lipid (red boxes) metabolism for *Rhogostoma schuessleri* (3EH3) based on KEGG ontologies and KEGG modules. Nodes represent components and edges represent enzymatic reactions. The edge color indicates the percentage of KO terms present, normalized to the minimum number of KO terms required for the respective reaction. Solid edges indicate that all KO terms were present for the respective reaction.

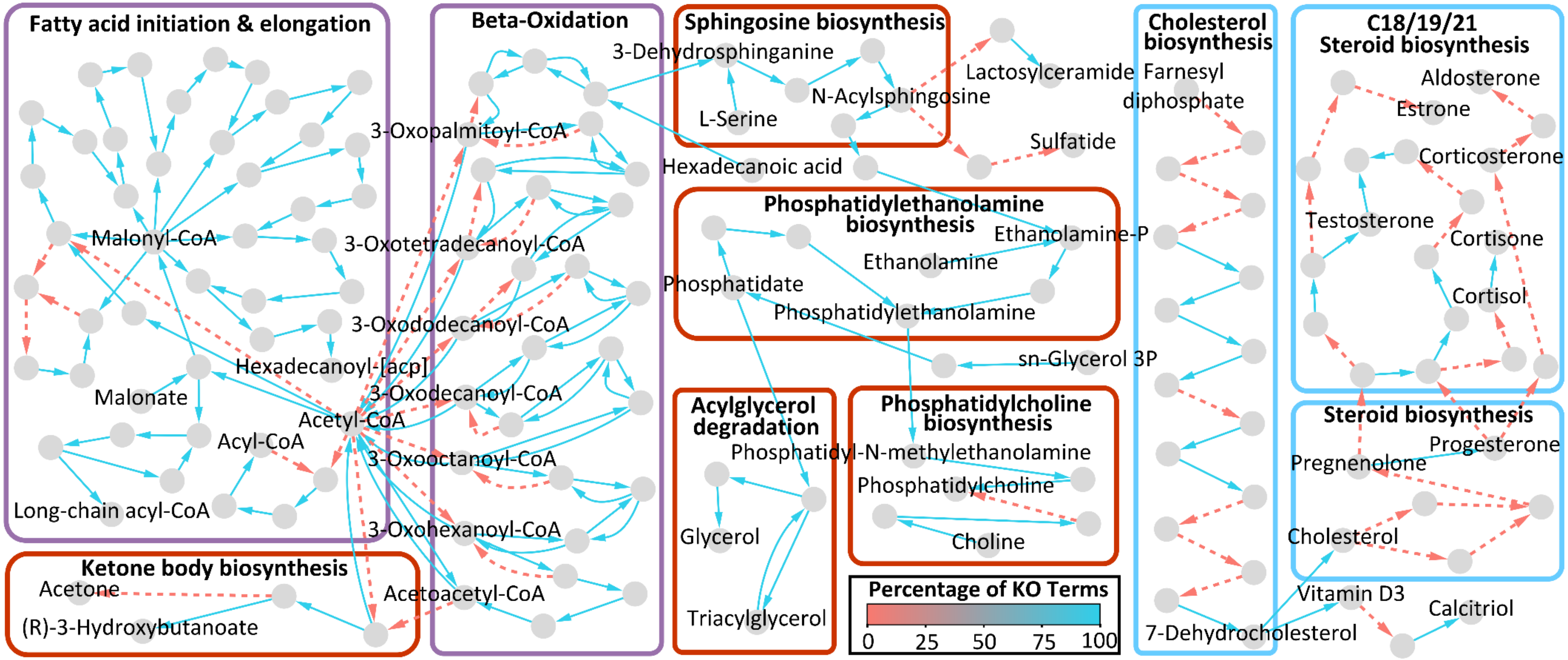


**Supplementary Figure 28: Overview of the lipid metabolism for *Fisculla terrestris*.** The graph illustrates a reconstruction of the fatty acid (purple boxes), sterol (blue boxes), and lipid (red boxes) metabolism for *Fisculla terrestris* based on KEGG ontologies and KEGG modules. Nodes represent components and edges represent enzymatic reactions. The edge color indicates the percentage of KO terms present, normalized to the minimum number of KO terms required for the respective reaction. Solid edges indicate that all KO terms were present for the respective reaction.

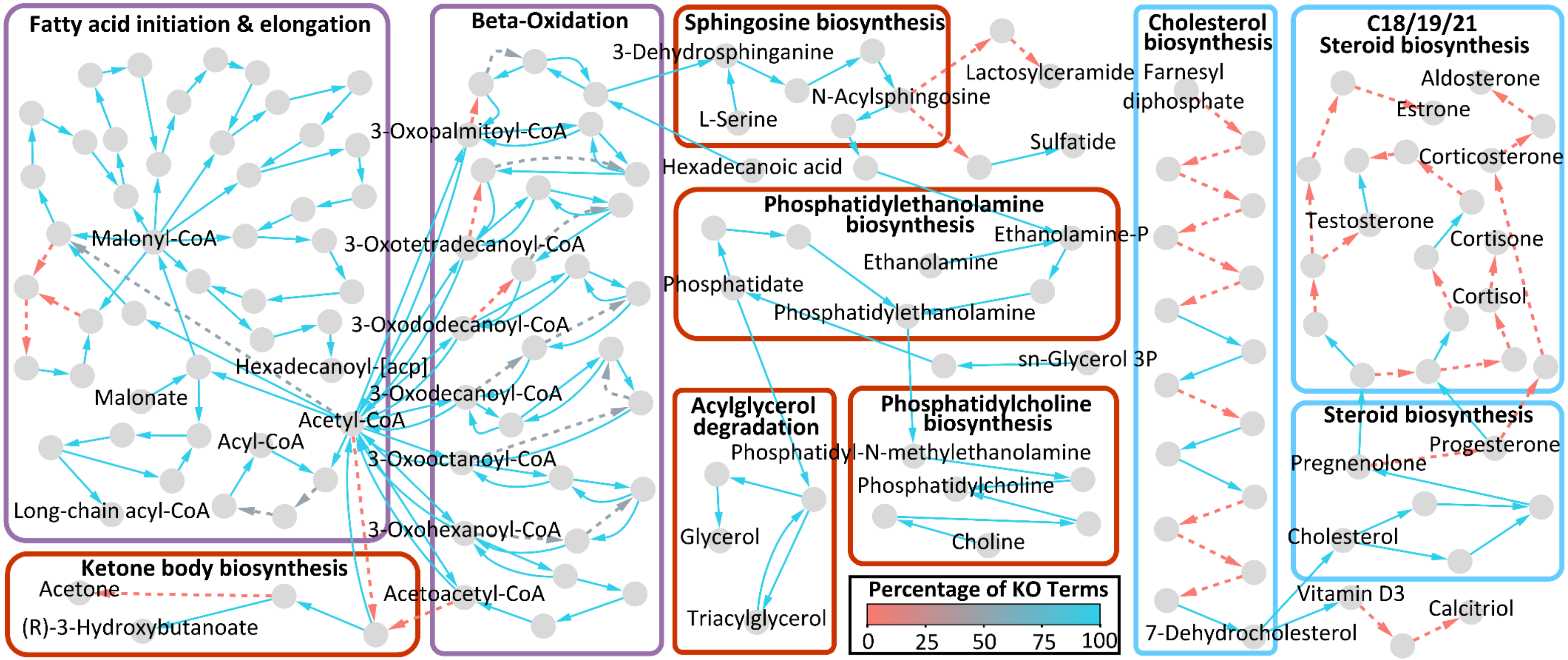


**Supplementary Figure 29: Overview of the lipid metabolism for *Katarium polorum*.** The graph illustrates a reconstruction of the fatty acid (purple boxes), sterol (blue boxes), and lipid (red boxes) metabolism for *Katarium polorum* based on KEGG ontologies and KEGG modules. Nodes represent components and edges represent enzymatic reactions. The edge color indicates the percentage of KO terms present, normalized to the minimum number of KO terms required for the respective reaction. Solid edges indicate that all KO terms were presentfor the respective reaction.
